# RNA transcribed from heterochromatic simple-tandem repeats are required for male fertility and histone-protamine exchange in *Drosophila melanogaster*

**DOI:** 10.1101/617175

**Authors:** Wilbur K Mills, Yuh Chwen G. Lee, Antje M Kochendoerfer, Elaine M Dunleavy, Gary H. Karpen

## Abstract

Long arrays of simple, tandemly repeated DNA sequences (known as satellites) are enriched in centromeres^1^ and pericentromeric regions^2^, and contribute to chromosome segregation and other heterochromatin functions^3,4^. Surprisingly, satellite DNAs are expressed in many multicellular eukaryotes, and their aberrant transcription may contribute to carcinogenesis and cellular toxicity^5-7^. Satellite transcription and/or RNAs may also promote centromere and heterochromatin activities ^8-12^. However, we lack direct evidence that satellite DNA transcripts are required for normal cell or organismal functions. Here, we show that satellite RNAs derived from AAGAG tandem repeats are transcribed in many cell types throughout *Drosophila melanogaster* development, enriched in neuronal tissues and testes, localized within heterochromatic regions, and important for viability. Strikingly, we find that AAGAG transcripts are necessary for male fertility and are specifically required for normal histone-protamine exchange and sperm chromatin organization. Since AAGAG RNA-dependent events happen late in spermatogenesis when the transcripts are not detected, we speculate that AAGAG RNA functions in primary spermatocytes to ‘prime’ post-meiosis steps in sperm maturation. In addition to demonstrating specific essential functions for AAGAG RNAs, comparisons between closely related *Drosophila* species suggest that satellite repeats and their transcription evolve quickly to generate new functions.

## Introduction

Simple, tandemly repeated satellite DNAs, such as AAGAG(n) and AATAT(n), comprise 15-20% of the *D. melanogaster* genome^13^. Given the emerging roles of non-protein coding RNAs (ncRNAs) in chromatin organization and other biological functions^14^, we investigated whether heterochromatic satellite transcripts are required for normal viability and development. We first analyzed RNA expression for 31 of the most abundant satellite DNAs, using published RNA-seq data (modENCODE)^15^ and RNA-Fluorescence In-Situ Hybridization (RNA-FISH) (Extended Data Fig. 1). Further characterizations and functional analyses were focused on AAGAG(n) RNA (hereafter AAGAG RNA) because it is highly abundant and linked to the nuclear matrix^16^.

## Results and Discussion

Northern blot analysis of RNA isolated from stage 1-4 embryos shows that AAGAG RNA is maternally loaded as an ∼1,500 nucleotide (nt) transcript. Smaller RNAs (∼20-750 nt) accumulate in later stage embryos (2-24hrs) and third instar larvae (L3 larvae) (Extended Data Fig. 2A). AAGAG RNA-FISH in 0-18hr embryos and L3 larvae revealed localization to only one or a few nuclear foci, with no visible cytoplasmic signal (Figs. 1A and B). AAGAG RNA foci are not detected prior to embryonic cycle 11, but by cycle 12, 33% of embryos have one or more foci and by cycle 13, 67% of embryos have one or more foci (Extended Data Figs. 2B and C). Furthermore, 100% of embryos exhibit nuclear AAGAG RNA foci by blastoderm (cycle 14, ∼2hrs after egg laying), coincident with the formation of stable, mature heterochromatin^17,18^ (Figure 1A and Extended Data Fig. 2D). Surprisingly, the complementary RNA (CUCUU(n)) is not observed in Northern or RNA-FISH analysis (Extended Data Fig. 4B and data not shown, respectively), suggesting that most or all of the AAGAG(n) DNA present at multiple genome locations are unidirectionally transcribed. This conclusion is supported by the results of RNase digestion experiments, which demonstrate that cycle 14 AAGAG RNA foci contain single-stranded RNA (ssRNA), and not R-loops or double-stranded RNA (dsRNA) (Extended Data Fig. 3). Finally, a combination of transcriptome mining, Northern blotting and RNA-FISH indicates that the majority of AAGAG RNA is transcribed from loci in 2R, X and 3R heterochromatin (Extended Data Fig. 4).

**Figure 1.**
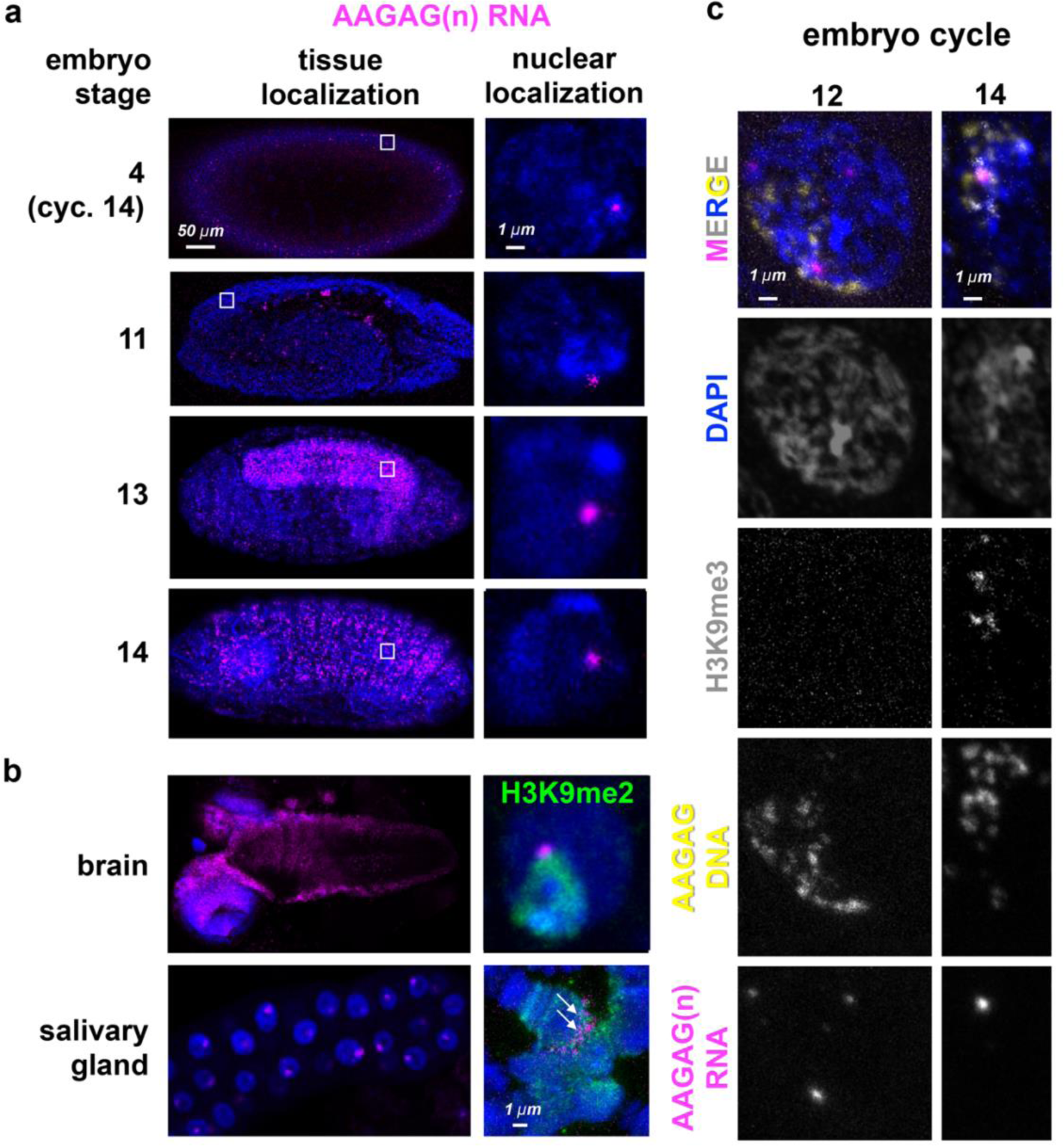
AAGAG(n) RNA localizations in embryos and larvae. **a**, AAGAG RNA distributions (magenta) throughout embryonic and larval development in Oregon R flies. DNA / DAPI = blue; all images are confocal sections. White box indicates location of enlarged nucleus (right side). **b**, Distributions of AAGAG RNA in intact larval L3 brain (top) and salivary gland (SG) tissue (bottom). Magnified images of projections of individual cells (right column) show that AAGAG RNA foci are located in or near the pericentromeric heterochromatin (H3K9me2 antibody IF, green) in brain cell nuclei. Note that foci are adjacent to nucleoli in intact salivary glands (SGs, bottom left), and at the chromocenter and not the euchromatic arms in the SG squash (arrows, bottom right). **c**, Projections of representative nuclei probed for AAGAG RNA (magenta) and AAGAG DNA (yellow),and stained for H3K9me3 (grey) and DNA (DAPI=blue). Left=cycle 12 nuclei prior to stable heterochromatin formation; right=early cycle 14 nucleus during heterochromatin formation. Note that in cycle 12, the few AAGAG RNA foci do not co-localize with AAGAG DNA. In cycle 14, AAGAG RNA foci co-localize with AAGAG DNA and H3K9me3.

**Figure 2.**
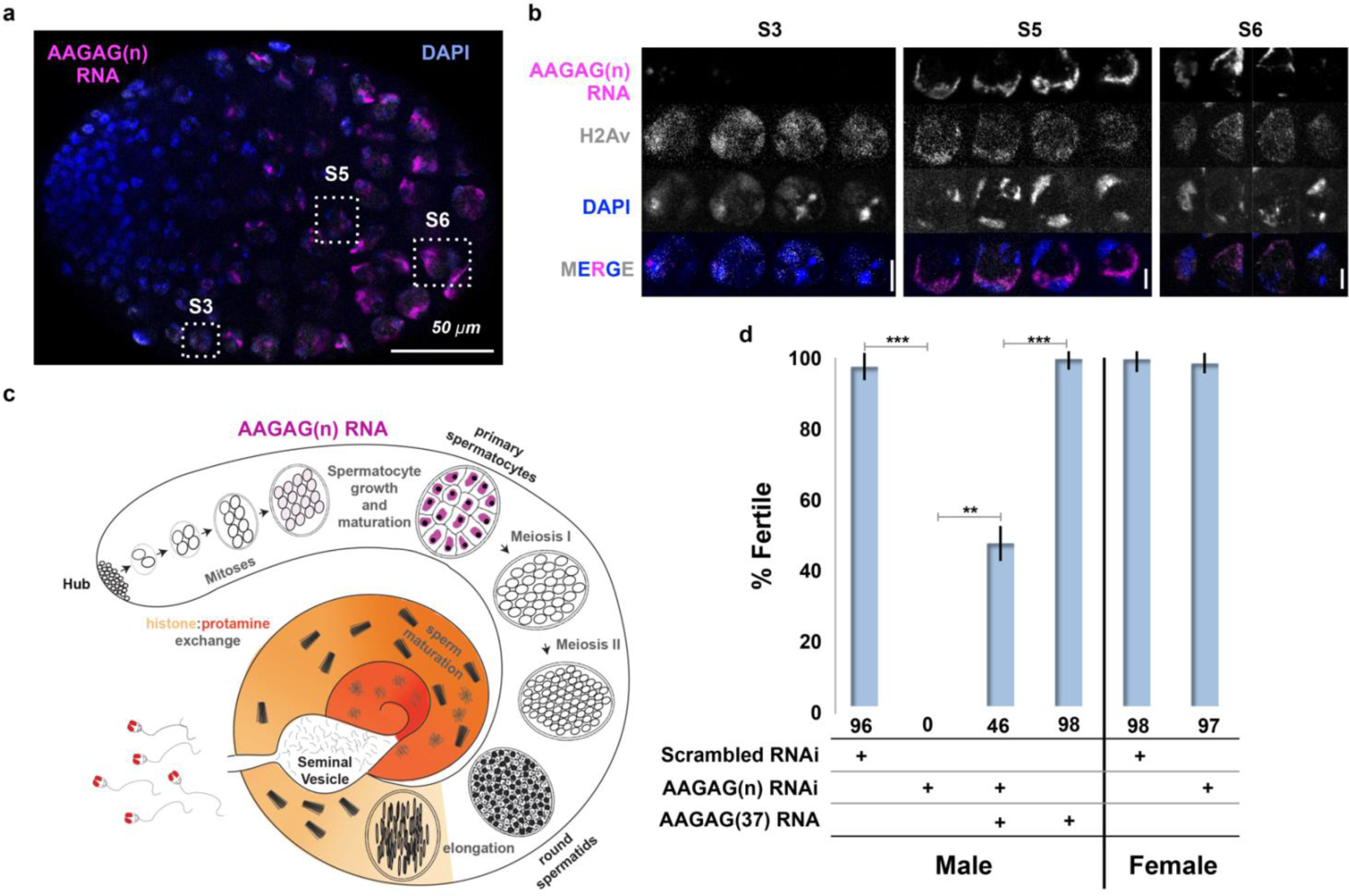
AAGAG RNA is enriched in primary spermatocytes and necessary for male fertility. **a**, Confocal section of a larval testis. RNA-FISH to AAGAG=magenta, H2Av (chromatin) IF=grey, DNA (DAPI)=blue. S3, S5, and S6 refer to primary spermatocyte stages. **b**, Enlarged confocal sections (representative boxes in a) of spermatocyte stages in larvae testes; scale bars=5µm. **c**, Schematic summary of AAGAG RNA (magenta) localization in adult testes (see Extended Data Figure 7 for a detailed description of spermatogenesis stages and events). AAGAG RNAs are visible in 16 cell primary spermatocytes (dark pink), and potentially 16 cell spermatogonial cysts (light pink); no AAGAG RNA was detected at earlier stages (hub, 2-8 cell spermatogonial cysts) or after the primary spermatocyte stage (meiosis I and II, sperm elongation and maturation). **d**, Fertility after depletion of AAGAG(n) RNA in male primary spermatocytes or female ovaries using the Bam-GAL4 driver. An ∼72% reduction in AAGAG RNA levels in testes (see Extended Figs. 9B and C) results in complete male sterility but has no effect on female fertility. Expression of AAGAG(37) RNA simultaneously with AAGAG RNAi (both driven by Bam-Gal4) partially rescues male sterility (46% fertile). Expression of AAGAG RNA alone, without depletion of endogenous AAGAG RNAs, has no impact on male fertility. Statistically significant differences based on T-tests are indicated by horizontal lines; *** p<0.001, **p<0.01,; variation is represented by stdev.

To determine where these transcripts localize within the nucleus, we simultaneously performed antibody staining (IF) for a histone post-translational modification enriched in heterochromatin (H3K9me3), and FISH for both AAGAG RNA and DNA. In cycle 12 embryos, AAGAG RNA is distributed randomly throughout the nucleus (Fig. 1C) and does not co-localize with AAGAG(n) DNA. Once stable heterochromatin forms (cycle 14)^18^, AAGAG RNA foci specifically co-localize with H3K9me3 (Fig. 1C). By stage 13 embryos (∼9.5 hrs after egg-laying) AAGAG RNA is specifically enriched in the ventral ganglia (neural tissue), and foci remain co-localized with heterochromatin (Fig. 1B). In addition, AAGAG RNA localizes to the chromocenter in polytene larval salivary glands (Fig. 1B).

The presence of AAGAG RNA throughout development suggested a potential role in development or viability. This hypothesis was tested by depleting AAGAG RNA in somatic cells, using actin-GAL4 driven AAGAG shRNA expression (Extended Data Fig. 6). Depletion of AAGAG RNA results in significantly lower viability by pupal stage compared to controls, with most lethality occurring during third instar larval (L3) stages (Extended Data Figs. 6G and H, respectively). We conclude that AAGAG RNA associates with the earliest forms of heterochromatin, maintains this localization throughout embryonic and larval development, is enriched in neural tissue and is important for viability.

Surviving act-GAL4 driven AAGAG RNAi adults exhibited partial sterility, prompting further investigation into the distribution and potential functions of AAGAG RNA in the germ line (see Extended Data Fig. 7 for an overview of spermatogenesis). In larval and adult testes, high levels of AAGAG RNA are observed in primary spermatocytes, where they are enriched in regions adjacent to DAPI-rich (AT-rich) DNA located at the nuclear periphery (Fig. 2A-C). AAGAG RNA is not detectable, even with amplified signal, at earlier stages near the hub, or at later stages (meiosis I and II, and subsequent stages of sperm development). Spermatocyte AAGAG RNA originates from the same 2R, 3R and X heterochromatic satellite regions identified in somatic cells and is specifically not from the Y chromosome (Extended Data Figs. 8A and B). To deplete AAGAG RNA we used the Bag of marbles (Bam)-GAL4^19^ driver to express AAGAG shRNA in 4-16 cell spermatogonial cysts. Strikingly, AAGAG depletion (∼72% reduction) results in 100% male sterility, with no impact on female fertility (Fig. 2D). AAGAG RNAi using drivers expressed earlier in spermatogenesis do not cause fertility defects (Extended Data Table 2). We conclude that expression of AAGAG RNA in primary spermatocytes is required for male fertility.

These results suggested that loss of male fertility after AAGAG RNA depletion would be caused by defects at stages where AAGAG RNA is expressed. Surprisingly, Bam-GAL4 driven depletion of AAGAG RNA resulted in no gross morphological defects prior to or during meiosis I or II in pupal or adult (0-6 hour and 4-7 days post-eclosion) testes (not shown). However, individualized mature sperm DNA was completely absent from the seminal vesicles (SV), in contrast to their abundance in controls (Fig. 3A), demonstrating that AAGAG RNA is important for later steps in spermatogenesis. In fact, the first visible defects are observed during individualization and maturation (Fig. 3B), stages that lack visible AAGAG RNA (Fig. 2C). For instance, aberrant individualization (irregular, long and decondensed sperm DNA) occurred at significantly higher frequencies after AAGAG RNA depletion, compared to scrambled RNAi controls (Fig. 3E). At later individualization stages, sperm bundles in AAGAG RNA depleted testes often contained less than the normal 64 sperm and were disorganized, displaying ‘lagging’ sperm nuclei and loosely packed sperm bundles (Fig. 3B and E). Finally, sperm DNA present was abnormally ‘kinked,’ ‘needle eyed’ or ‘knotted’ in appearance, and normal, mature forms of sperm DNA readily found in basal regions (just prior to entry into the seminal vesicle) of control testes were never observed after AAGAG depletion (Fig. 3B). These phenotypes indicated that AAGAG RNA is important for sperm nuclear organization, and were similar to the consequences of defective histone-protamine transitions observed previously^20,21^. Strikingly, antibody IF revealed that Bam-GAL4 driven AAGAG RNA depletion caused reduced and defective incorporation of the transition protein Mst77F (Fig. 3C), and an absence of Protamine A/B (Fig. 3D). We conclude that AAGAG RNA is required for fertility at least in part by promoting the histone-protamine transition.

**Figure 3.**
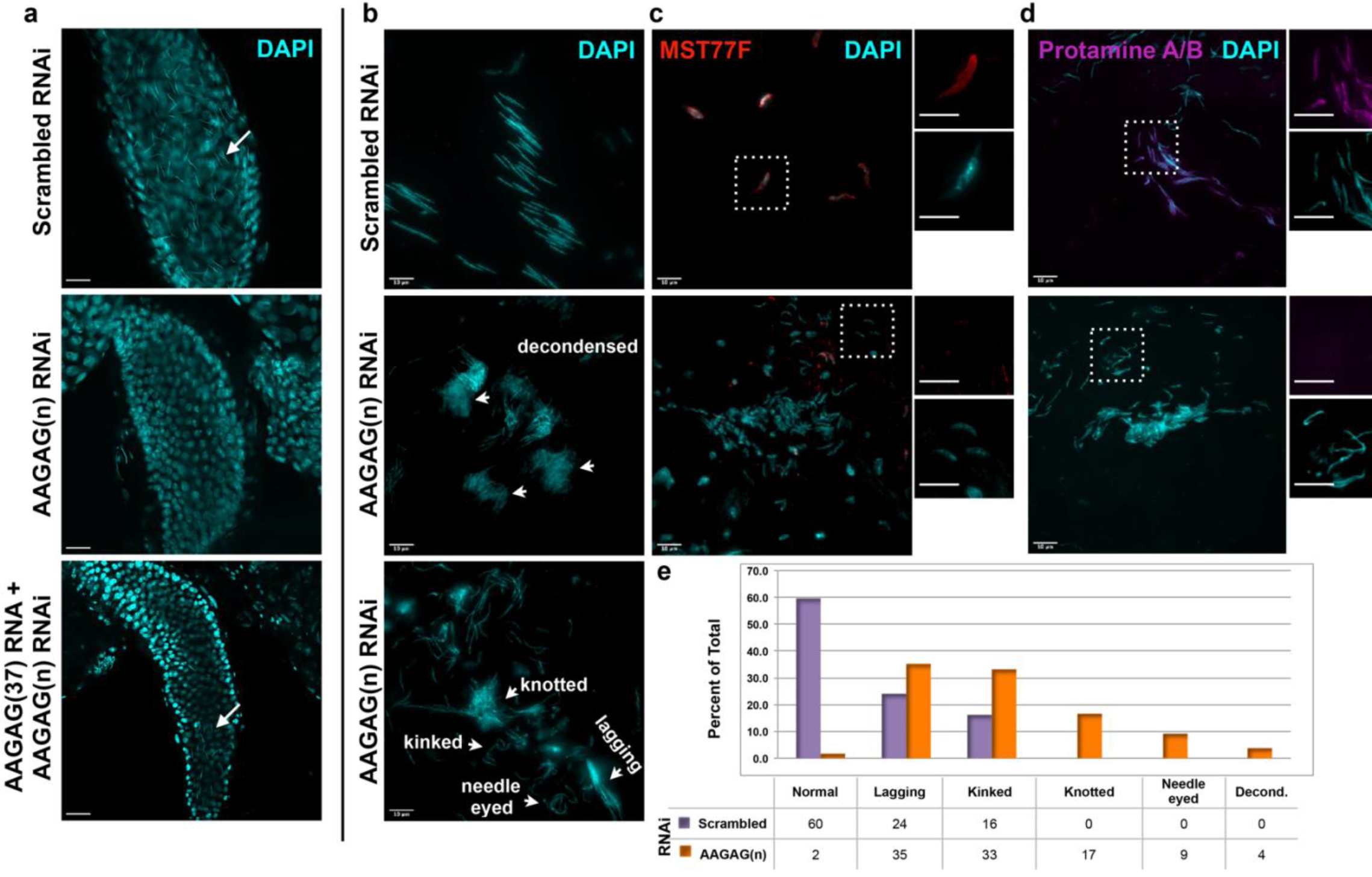
AAGAG RNAi depletion in mitotic germline cysts and spermatocytes (Bam-GAL4 driver) results in severe defects in sperm maturation and protamine deposition. **a**, Seminal vesicles (SVs) in testes from 0-6 hour old adults; DAPI (DNA)=cyan. Mature sperm nuclei visible as thin, elongated DAPI signals in the scrambled control (top, white arrow) are absent after AAGAG RNAi. Individualized mature sperm (white arrow) are visible in SVs from AAGAG RNAi males that also express AAGAG(37) RNA (partial rescue, 4-7 day old adults). **b**, Bundles of elongating sperm nuclei visible in the scrambled RNAi control (top). Defective ‘decondensed’ (middle, white arrowheads), ‘knotted,’ ‘kinked ‘needle eyed’ and ‘lagging’ (bottom, white arrowheads) sperm phenotypes are visible in the AAGAG RNAi but are much less frequent or absent in controls (see **e**). **c**, Transition Protein Mst77F (red) is present on sperm in control RNAi but is largely absent and/or disorganized after AAGAG RNAi (dashed boxes indicate regions in the zoomed images to the right). **d**, Protamine A/B (purple) is present on sperm in the scrambled control RNAi but is absent after AAGAG RNAi. Scale bars = 10 µm except for zoomed images in c and d = 8 µm. **e**, Quantitation of sperm defects associated with RNAi depletion of AAGAG RNA in 4-6 day old adults, compared to scrambled RNAi control.

Importantly, fertility defects resulting from AAGAG RNA depletion are partially rescued by simultaneously expressing AAGAG RNA (185 bases, 37 repeats), when both are controlled by the Bam-GAL4 driver. Under these conditions we observe a 2-fold increase in AAGAG RNA signal compared to AAGAG RNAi alone (Extended Data Fig. 9C), which is sufficient to restore male fertility (46% with AAGAG RNA expression compared to 0% in AAGAG RNAi alone, Fig. 2D) and the presence of mature sperm in the seminal vesicles (Fig. 3A). Together, these results demonstrate that RNA from the simple tandem repeat AAGAG(n) is necessary for spermatogenesis in *Drosophila melanogaster*.

Here, we demonstrate that AAGAG(n) satellite RNAs are transcribed from heterochromatic regions on multiple chromosomes, cluster into one nuclear focus, associate with the earliest forms of heterochromatin in embryos, persist throughout fly development and are important for viability. Most importantly, AAGAG RNA is necessary for male fertility, and is specifically required for mature sperm formation and successful histone-protamine exchange. Notably, AAGAG RNA is present prior to meiosis, yet is required for processes in later spermatogenesis stages, when the RNA is not detected. We speculate that AAGAG RNA ‘primes’ primary spermatocyte nuclei to successfully complete post-meiotic sperm development. AAGAG RNA could sequester or exclude factors that need to be localized to specific spermatocyte regions or loci (Figs. 4A and B) or could affect formation of essential complexes (Fig. 4C), levels of RNA, proteins, or post-translational modifications, and/or organization of chromatin in spermatocytes. Regardless, it is surprising that AAGAG RNA in primary spermatocytes is necessary to promote critical sperm development at much later stages, including but not limited to histone-protamine exchange (Figs. 4D and E).

**Figure 4.**
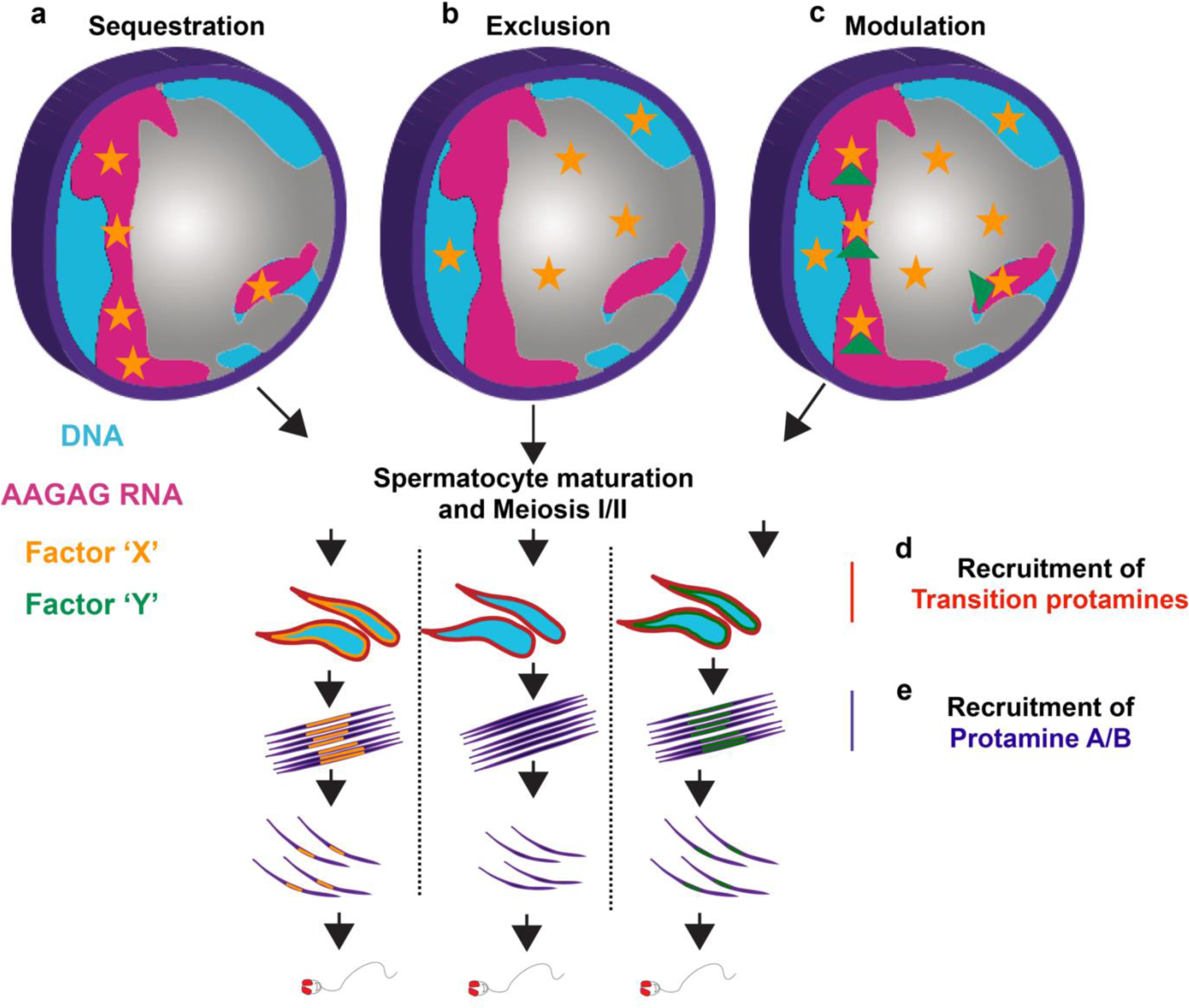
Model for AAGAG RNA function during spermatogenesis. **a**, AAGAG RNA may sequester factors (orange star, i.e. RNA/DNA/proteins) or **b**, exclude factors from specific locations within primary spermatocyte nuclei. **c**, AAGAG RNA may also affect formation of complexes (orange star plus green triangle), or genome organization, expression levels, or post translational modifications (PTMs) in primary spermatocytes. Based on the defects observed in late stages of spermatogenesis after AAGAG RNA depletion, such as the histone-protamine transition, activities promoted by the presence of AAGAG RNA in primary spermatocytes are not required until later stages, including **d**, recruitment of transition protamines such as Mst77F (red) in leaf and canoe stage spermatids, **e**, recruitment of protamines (purple) in sperm bundles, and/or unknown factors required for mature sperm formation.

Finally, it is worth noting that the expression of simple satellites for essential functions seems incompatible with the fast evolution of satellite DNAs, reflected in dramatic changes in both sequence types and copy numbers across species^22^. Specifically, AAGAG is one of the most abundant simple repeats in *D. melanogaster*, comprising ∼5.6% of the genome^23^. However, the amount of AAGAG is several orders of magnitude lower in the closely related *D. simulans* and *D. sechellia*, and is nearly absent in other *Drosophila* species^22^. It is possible that in species with few or no AAGAG repeats, low levels of AAGAG RNA are sufficient for fertility, or the expression of different lineage-specific satellite arrays are required for protamine exchange and other events in sperm maturation. The fast turnover of AAGAG DNA between species and its critical functions in male reproduction mirror the frequently observed adaptive evolution of new lineage-specific protein-coding genes^24^, which are biased toward testis-expression and acquisition of essential functions in male reproduction, including spermatogenesis^25^. Selective pressures proposed to drive the fast evolution of new testis-expressed genes^26^, such as sperm competition, sexual conflict, antagonistic interactions with germline parasites and/or selfish DNAs, could also drive the fast turnover and acquisition of essential functions by non-coding satellite DNAs. Thus, our results provide a strong impetus for additional studies of satellite RNA functions, which could elucidate new roles of so-called “junk DNA” in health, disease and evolution.

## Acknowledgements

WKM and GK conceived of and designed the project, and wrote the manuscript. EMD provided critical guidance in spermatogenesis experiments. YCGL identified the genomic sources of AAGAG transcripts. AMK performed histone-protamine transition staining in testes and WKM performed all other experiments. All authors provided critical edits and advice. We thank Karen Miga and Miten Jain for RNA-seq analysis efforts, Shelby Wilson for technical guidance with spermatogenesis analysis, and Aniek Jansen for editing.

## Funding

National Science Foundation Graduate Research Fellowship to W.K.M. (ID:2013160110)

NIH RO1 # GM117420 to G.K.

NIH K99 GM121868 to YCGL

## Materials and Methods

### Imaging

Most images were acquired using a Zeiss LSM710 confocal microscope using 40X water or 63X oil objectives. For these confocal images, projections were acquired as z-stacks with step sizes depending on the sample. Image files were then processed and analyzed using Fiji. Non-rescue testes images in Fig. 3 were acquired using DeltaVision Elite wide-field microscope system (Applied Precision). Images were acquired as z-stacks with a step size of 0.5 μm, raw data files were deconvolved using a maximum intensity algorithm. 3D z-stack images were represented in 2D by projection using SoftWorx (Applied Precision).

### RNA probe generation for RNA-FISH

RNA probes were made by using oligo templates with antisense T3 promoters on the 3’ends, hybridizing an oligo composed of sense T3 promoter so as to create a double stranded 3’ end, or in the case of 359 bp repeat, amplification with oligos containing T3 and T7 promoter ends on genomic DNA using standard protocols. Probe templates were then transcribed with T3 RNA polymerase (or T7 for one strand of 359 bp repeat) and either UTP-biotin or UTP-digoxeginin labels, or in the case of RNA without Uracil, biotin-ATP. Oligos are listed in Table 1 and were ordered standard desalted from IDT. Reaction conditions were as follows: In a 40ul reaction, 1X RNApol reaction buffer (NEB cat. MO3782), 1mM each final concentration of ATP, GTP, CTP and 0.62 mM UTP, supplemented with 0.35mM final concentration of either digoxegenin-11-UTP (Roche cat. 3359247910), biotin-UTP (Sigma, cat. 11388908910), or biotin-11-ATP (Perkin Elmer, cat. NEL544001EA), 1 Unit Protector Rnase inhibitor (Roche cat. 3335402001), 5µM each of probe template and T3 promoter oligo (5’-AATTAACCCTCACTAAAG), and H_2_0 to 40µl were combined. Reactions were heated to 80°C, 3min to denature probes, iced 2 min., 4µl (or 200Units) of T3 (or T7) RNA polymerase (NEB cat. M0378S) added and incubated at 37°C overnight. 2µl Turbo DNAse (ThermoFisher Cat. AM2238) was then added to degrade DNA templates, incubated at 37°C for 15 min and the reaction stopped by adding 1.6µl of 500mM EDTA. Probes were then purified using standard sodium acetate/ethanol purification. Probe concentration was then assessed using Qubit RNA high sensitivity protocols and reagents and stored at −80°C.

**Table 1.**
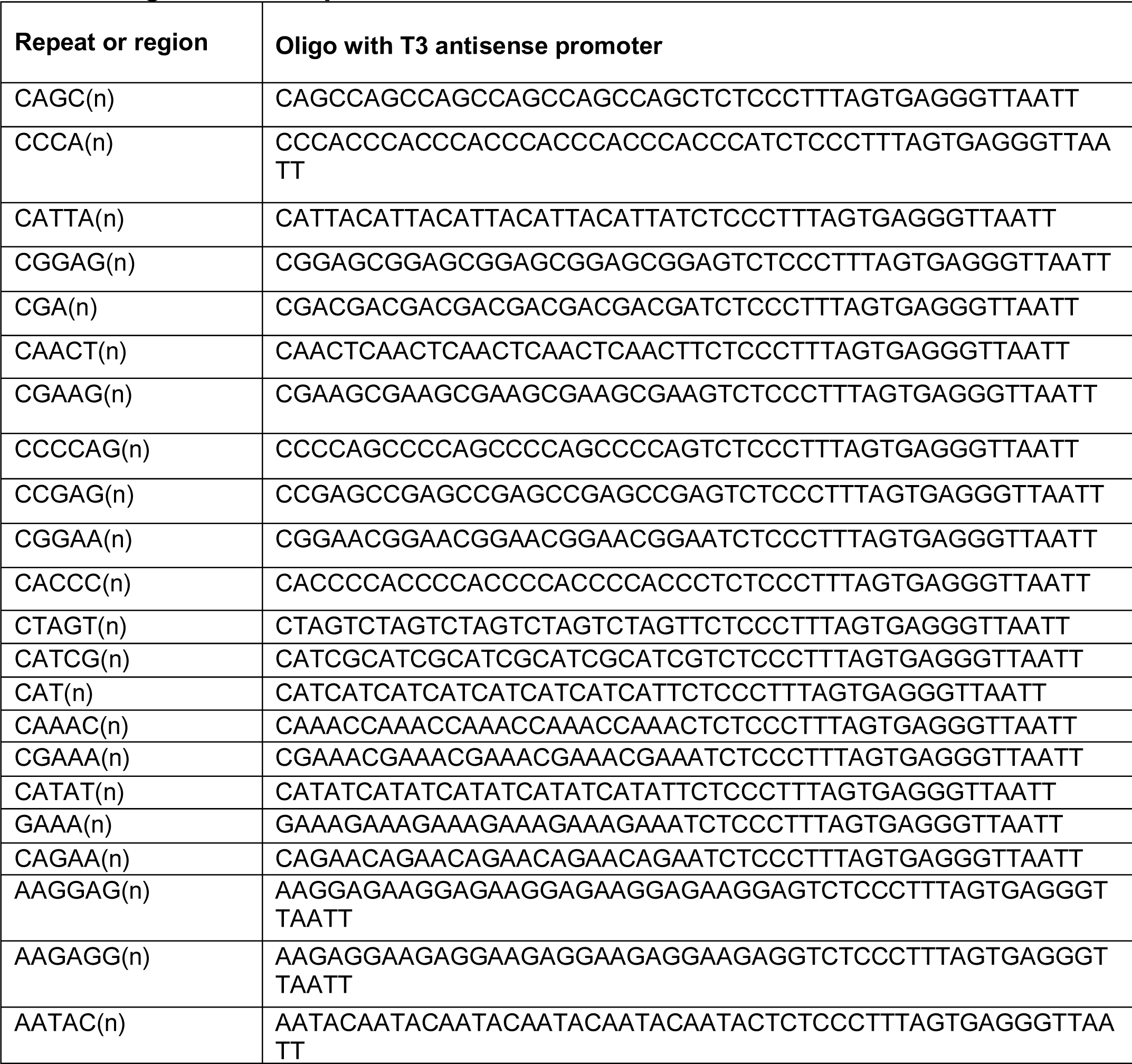
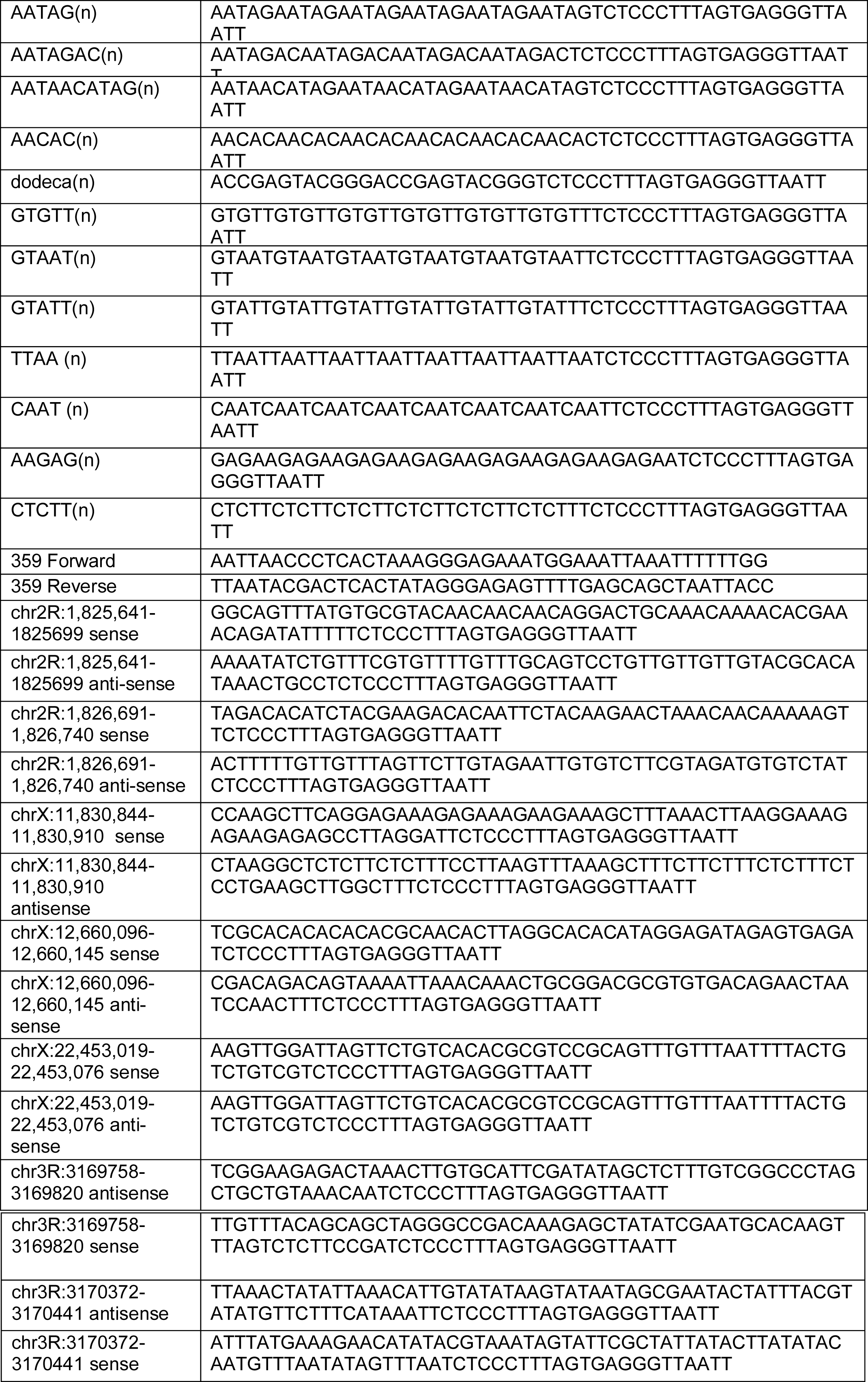
Oligos for RNA probes.

### RNA-FISH Buffers

<u>PBT solution</u>: 1X PBS and 0.1% Tween-20; <u>Western Blocking Reagent 10X:</u> 10% casein in 100mM maleic acid; 150mM NaCl; pH 7.5. heated at 60°C for 1hr to dissolve. <u>PBT block</u>: 1:1 PBT/2X WBR; <u>Hybridization buffer</u>: 50% formamide, 5X SSC, 100 μg/mL heparin, 100 μg/mL sonicated salmon sperm DNA, and 0.1% Tween-20, filtered through a 0.2μm filter.

### For clarity, the methods for RNA-FISH probe hybridization and detection are numbered below

#### RNA-FISH methods

##### Protocol 1. RNA-FISH probe hybridization and primary antibody incubation

RNA probe hybridization for all tissues was carried about according to^27^, steps 10-17 under subheading #3. Samples were then washed one time with PBT then blocked in PBT block 1hr at room temperature. Samples were then processed for either ‘non-Tyramide Signal Amplification (TSA) probe amplification’ (Protocol 2) or ‘TSA amplification for RNA-FISH probe detection” (Protocol 3)

##### Protocol 2. Non-TSA probe detection for RNA-FISH

For ‘non-TSA amplification’, samples were incubated with either mouse anti-digoxigenin-Cy5 (source unknown) or rabbit anti-digoxigenin A488 (Invitrogen cat# 700772) in PBT block at 1/200 dilution for 1hr at room temperature. Afterwards, samples were incubated 6x’s 10 min each in PBT block, stained with DAPI 10min, washed 3’xs 10min. each in PBS and mounted in Prolong-Gold antifade mountant (Thermofisher, cat. P36390). **Figures processed using this protocol:** Fig 1B, right panel; Figs.2 A and B; Extended Data Figs. 4C-H for AAGAG only; Extended Data Figs. 6B-D; Extended data Figs. 8 and 9.

##### Protocol 3. TSA amplification for RNA-FISH probe detection

For samples undergoing ‘TSA amplification for RNA-FISH probe detection,’ samples were incubated with primary antibody (1/400 dilution of mouse anti-digoxegenin coupled to biotin (Jackson Immuno Research cat.200-062-156, lot. 123482)), with 0.2 U/µl protector RNAse inhibitor and incubated overnight at 4°C. Next, samples were washed 6x’s 10min each in PBT block. The next steps are essentially as per ‘tyramide signal amplification kit’ protocols (ThermoFisher) but with reagents purchased separately: Samples were incubated with 1:100 streptavidin-HRP (Molecular probes, cat. S911) in PBT block for 1hour at room temperature. Samples were then washed in 1:1 PBT/2XWBR 6x’s 10min each, once with PBT, and 2x’s with PBS. Samples were then incubated with Alexa 647 tyramide (TSA™ Reagent, Alexa Fluor® 647 Tyramide cat. T20951) according to company protocols. Essentially, this consisted of adding 1µl of 30% hydrogen peroxide to 200µl tyramide signal kit amplification buffer, then diluting this solution 1/100 in tyramide signal amplification buffer for a final hydrogen peroxide concentration of 0.0015%. This solution was then added to the sample and incubated at room temperature for 1hr in the dark. Samples were then washed 1x with PBS for 10min, stained with DAPI for 10min, washed 4x’s with PBS 10min. each, and mounted in Prolong Gold Antifade mountant. **Figures processed using this protocol:** Figs 1A, both panels, left panel in 1B and both panels in 1C; Extended data Figs. 1-3; Extended data Figs. 4C-H for non AAGAG RNA detection (ie 2R and X heterochromatic transcripts); Extended Data Fig. 5.

### RNA-FISH of repeats in embryos

For RNA-FISH of repeat RNAs, 0-8hr Oregon R embryos were collected on apple juice plates, dechorionated and processed according to ^27^, as per protocols 1 and 3 above, with the exception of using 37% formaldehyde stock from Sigma (cat. F1635-500ML).

### Co-IF DNA/RNA-FISH of AAGAG RNA in embryos

(Fig. 1C). Co-IF RNA/DNA-FISH was performed essentially as described in ^28^, in which RNA-FISH was performed first, signal detected via tyramide signal amplification, RNAse treatment to remove RNA and prevent DNA-FISH probes binding to RNA, and then DNA-FISH performed. Essentially, RNA-FISH was performed as above, but after tyramide signal amplification (protocols 1 and 3 above) and washing, samples were fixed in 4% formaldehyde. Samples were then washed 3x’s in PBS 2min. each. RNA was then removed under the following conditions: In a 50µl final volume, 1X Shortcut RNaseIII buffer (NEB cat. M0245S), 1.5ul RNASEIII (neb cat. MO245S), 100µg/ml RNaseA final concentration, 1X MnCl2 (NEB cat. MO245S) and water to 50µl were added and samples incubated overnight at 4°C. Samples were then rinsed 3x’s in PBT 5min each, rinsed in 1:5, 1:1 and 5:1 mixtures of PBT: RNA hybridization solution for 15min each. Samples were then replaced with hybridization buffer and incubated 15 min. A DNA oligo probe to AAGAG(7) tagged with alexa5 was then diluted in hybridization buffer to 2.5ng/µl, denatured at 70°C for 3min, then left on ice for 2min. Hybridization solution was removed from the embryos, probe solution added, and the sample denatured at 80°C for 15min and hybridized overnight at 37°C with nutation. Samples were then washed 2x’s with pre-warmed 37°C hybridization buffer 10 min each. Samples were then washed in 3:1, 1:1, 1:3 hybridization buffer:PBT 15min each at 37°C. Samples were then washed 2x’s in PBT at room temperature 5 min each. Samples were then stained with DAPI 10min, washed once in PBS, and mounted in Pro-Long Gold Antifade mountant.

### RNA-FISH in larvae

This protocol is essentially as described in ^29^. All figures containing larval RNA-FISH (Fig 1B left column and top right nuclei, Figs. 2A and B and extended data Fig 4C-H) used protocol A) and C) below. Those processed for TSA (needed for protocol 3 above) additionally used B below.

A. Third instar larvae were dissected in PBS supplemented with 0.2U/µl Protector RNase Inhibitor. The posterior end of the larvae was removed, then the remaining L3 inverted inside out. The inverted larvae were then transferred to ice cold PBS with 0.2U/µl RNAse inhibitor. Larvae were then fixed in PBT with 4% formaldehyde for 15min, then washed 3x’s, 5min each with PBT. Larvae were then incubated with 0.1%(vol/vol) DEPC in PBT for 5min to deactivate endogenous RNAses. Samples were then rinsed 2x’s with PBS.
B. Use of TSA amplification in L3 requires removal of endogenous peroxidases and requires the following protocol after DEPC treatment above and rinsing in PBS: In order to quench endogenous peroxidases, samples were incubated in 350μl (enough to cover all tissue) of 3% H_2_O_2_ in PBS 15 min at room temperature and the tube kept open to prevent gas buildup. Samples were then rinsed 2x’s with PBT 10min. each.
C. To all larval samples: Larvae were then permeabilized by incubation in 500µl cold 80% acetone in water at −20°C 10min. Samples were then washed 2x’s, 5min. per wash with PBT, then post fixed with 4% formaldehyde in PBT for 5min. Samples were then washed 5x’s with PBT 2min each. Samples were then rinsed with 1:1 PBT/RNA hybridization solution, then with 100% RNA hybridization solution, and then stored in hybridization solution at −20°C until needed. Samples were then processed according to RNA-FISH protocol (protocol 1 above, under ‘RNA FISH methods’) for probe hybridization and either (protocol 2 above, under ‘RNA FISH methods’) for non-TSA probe or (protocol 3 above, under ‘RNA FISH methods’) for TSA amplification.

### RNA-FISH in salivary gland squash

(Fig 1B, bottom right) Larvae were grown, prepped and salivary glands processed as per ^30^, rehydrated in 95%, 70%, then 30% ethanol 1min each, then washed 5min in PBT (0.1% Triton X-100 (TX100)). Slides were then fixed again in 3.7% formaldehyde in PBT (0.1% TX100), washed 2x’s 3min. each in PBT (0.1% TX100), treated with 0.1% DEPC in PBT (0.1% TX100) and washed one time in PBT (0.1% TX100). Sample was then covered with pre-denatured hybridization solution, covered with a coverslip and incubated at 56°C in a sealed hybridization chamber for 2 hours. The probe solution was then created by adding 100ng probe in 100µl hybridization solution, heating at 80°C for 3min., and cooling on ice for 5min. This probe solution was then added to the sample, a coverslip added and sealed with rubber cement, and incubated overnight at 56°C in a humid box. At 55°C in a coplin jar, slides were then treated in 50% formamide/PBT (0.1% tx100) 1hr, 25% formamide/PBT (0.1% Tx100) 10min, then 3x’s with PBT (0.1%Tx100) 10min each. Once at room temp, samples were blocked in 1:1 PBT/2xWBR and processed as per larval RNA-FISH using non -TSA probe detection (protocol 2 above).

### RNAse treatment of embryo

(Extended Data Fig 3). For RNAse of embryos prior to probe hybridization: RNA-FISH to AAGAG was performed on embryos pre-treated with RNaseIII (which cleaves dsRNA^31^), RNaseH (which cleaves the RNA strand in RNA/DNA hybrids), RNase I (which non-specifically cleaves ssRNA and dsRNA), and RNaseA (which cleaves adjacent to pyrimidines, preferentially in ssRNA, and specifically not between purines such as 5’-AGAAGGGAGAAG^32,33^). Reaction conditions were as follows: Samples were treated in 50µl final volume with either RNAseIII: 1X RNAseIII buffer, 1.5µl Shortcut RNaseIII (New England Biolabs, cat. M0245s), and 1X MnCL_2_; RNAseH treatment: 1X RNAseH buffer, 1.5µl RNAseH (New England Biolabs, cat. M0297S); RNAse 1 treatment: 1X RNAseH buffer, 1.5µl RNAse1 (Ambion cat. AM2294); RNAse A treatment: 1x RNAseH buffer, 15µg RNAseA-at 37°C for 5 hours. Samples were then washed 5x’s in PBT 2min each, treated with 0.1% DEPC to deactivate any remaining RNAse, then washed in PBT. Samples were then rinsed in 1:1 mixture of PBT:RNA hybridization solution for 2 min and resuspended in 100% hybridization solution. Samples were then processed as per ‘RNA-FISH probe hybridization and primary antibody incubation’ (protocol 1 above) and protocol ‘TSA amplification for RNA-FISH probe detection’ (protocol 3 above).

### RNAse of embryos after probe hybridization

(Extended Data Fig. 5). After probe hybridization and washing with PBS, samples were treated in 50ul final volume for either RNAseIII treatment: 1X RNAseIII buffer, 1.5µl Shortcut RNaseIII (New England Biolabs, cat. M0245s), and 1X MnCL_2_ or RNAseH treatment: (1X RNAseH buffer, 1.5µl RNAseH (New England Biolabs, cat. M0297S) at 37°C for 2 hours. Samples were then blocked with 2x PBT:WBR 1hr then processed as per protocol ‘TSA amplification for RNA-FISH probe detection’ (protocol 3 above).

### RNA-FISH in adult testes

For analysis of AAGAG RNA in RNAi adult testes (Extended Data Figure 9), flies were mated at 29°C and F_1_ progeny grown at 29°C to mimic conditions used to assess sperm morphological defects. AAGAG RNA was also visualized in RNAi testes grown at 25°C to rule out that temperature affected levels and distribution of AAGAG RNA (not shown). For analysis of AAGAG RNA in Oregon R and XO/XY testes (Extended Data Fig 8), flies were grown at 25°C. Flies were then anaesthetized with CO_2_, testes removed with forceps and placed in 7µl of PBS on (+) charged slides, a RainX-treated coverslip placed over the testes and both snap frozen in LiN_2_. The coverslip was then immediately removed with a razor blade and slides stored at −80°C until needed. When ready to process, slides were fixed for 20min in 4% formaldehyde in PBT, washed three times, 5min. each wash, in PBT. Samples were then incubated in 80% cold acetone in PBT for 10min at −20°C and processed as per RNA-FISH for ‘all larval samples’ using protocol 2 above for detection without TSA amplification.

### Northern blotting

Non-radioactive, denaturing northern blots were essentially carried out according to Chemiluminescent Nucleic Acid Detection Module Kit (Thermofisher cat# 89880). Essentially, purified RNA was denatured for 3 min at 70°C in NorthernMax formaldehyde loading dye. Samples were then run on denaturing agarose gels with 6.9% formaldehyde in MOPS buffer. RNA was transferred to (+) charged nylon membranes in an electroblotter (FisherBiotech Semi-Dry blotting unit, FB-SDB-2020) using 200mA for 30min. The membrane was then UVC crosslinked and prehybridized with ULTRAhyb Ultrasensitive Hybridization Buffer (Thermofisher, cat# 8669) at 68°C for 30min. Biotinylated probes at a concentration of 30ng/ml were then added to UltraHyb buffer, pre-hybridization solution replaced with solution containing probe and hybridized overnight at 68°C. The next day, membranes were washed and processed according to Chemiluminescent Nuclei Acid Detection kit manual.

### Identification of source of AAGAG RNA

To identify the genomic origin of AAGAG RNA, we mined *D. melanogaster* transcriptome data (modENCODE staged embryo and L3 larvae total RNA-seq reads)^15^ for AAGAG RNA attached to mappable ends with uniquely mapped sequences and focused on regions adjacent to >50bp blocks of annotated AAGAG(n) DNA. More specifically, we first used trim_galore to filter out adaptors and low quality sequencing reads. Reads with at least three consecutive AAGAG repeats were identified and their corresponding pair-end sequences were extracted. Including only AAGAG containing reads, we used Phrap to assemble the other end sequences into contigs using Phrap (-vector_bound 0 -forcelevel 5 -minscore 30 -minmatch 10). We then Blasted (with e-value < 10^−5^) to identify potential genomic locations in release 6 of *D. melanogaster* genome^34^ (Table 2). This conservative analysis revealed that the majority of AAGAG RNA originates from 2R and X heterochromatic satellites (Table 2 and Extended Data Fig 4). To confirm that this computational genomic analysis identified sources of AAGAG transcripts, we performed northern blotting and RNA-FISH to these and a 3R heterochromatic region. Essentially, transcript sizes using probes to these regions are similar if not identical to AAGAG RNA, and foci from these mappable regions co-localize with AAGAG RNA foci (Extended Data Figs. 4B and C-H, respectively), demonstrating that AAGAG RNA originates from identified 2R, X and 3R heterochromatin genomic regions.

**Table 2.** Uniquely mapped RNA identified via phrap adjacent AAGAG(>10) containing blocks.

| chr | e0-2hr | e2-4hr | e4-8hr | e8-12hr | e12-14hr | e14-16hr | e16-20hr | e20-24hr |
| --- | --- | --- | --- | --- | --- | --- | --- | --- |
| 2R | NA | NA | NA | NA | NA | NA | chr2R.18256<br>40.1825699 | NA |
| X | NA | NA | NA | NA | NA | NA | NA | chrX.126600<br>77.12660134 |
| X | NA | NA | NA | NA | NA | NA | chrX.118307<br>95.11830858 | NA |
| X | chrX.2245<br>3019.2245<br>3120 | NA | chrX.224<br>53019.22<br>453182 | NA | chrX.224<br>53019.22<br>453163 | chrX.224530<br>19.22453177 | chrX.224530<br>19.22453093 | chrX.224530<br>19.22453196 |

### Cloning of shRNA and over-expression constructs into pValium20 vector

Essentially, sense and antisense strands were annealed together, then ligated into digested pValium20 vector. For annealing, in a 50µl final volume, 1.5µl each of 100µM stock oligos were added to 1X NEBuffer, incubated 4min at 95°C, then placed in and slowly cooled (which was critical) in a 1L beaker filled with 70°C water. Once the water cooled to room temperature, samples were made blunt ended with klenow using standard procedures, purified with min-elute PCR purification kit, run on agarose gel, and appropriate size bands removed and purified. Purified bands were then digested with Nhe1 and EcoR1 HF enzymes and purified with min-elute PCR purification kit. For cloning, 1µl of annealed and purified oligo pair complement was added to 30ng of digested pValium20 vector and ligated with T4 DNA ligase (importantly not quick ligase) at 16°C overnight and transformed into dh5alpha *E. coli* cells for propagation.

### Insertion of shRNA or overexpression constructs

RNAi and overexpression lines were created via small-hairpin RNA (shRNA) to AAGAG RNA driven with the UAS/GAL4 system, or in the case of controls, a scrambled RNA sequence as well as mCherry, using genomic insertion of the pValium20 vector used for the Transgenic RNAi project (TRiP) at Harvard^35^. Importantly, the scrambled shRNA sequence contained the same percentage of A’s and G’s as in the test shRNA, but in a random order, to identify phenotypes resulting specifically from AAGAG knockdown versus off-target affects. pValium20 constructs were injected and screened for insertion by Rainbow Transgenic, Inc.

**Table 3.**
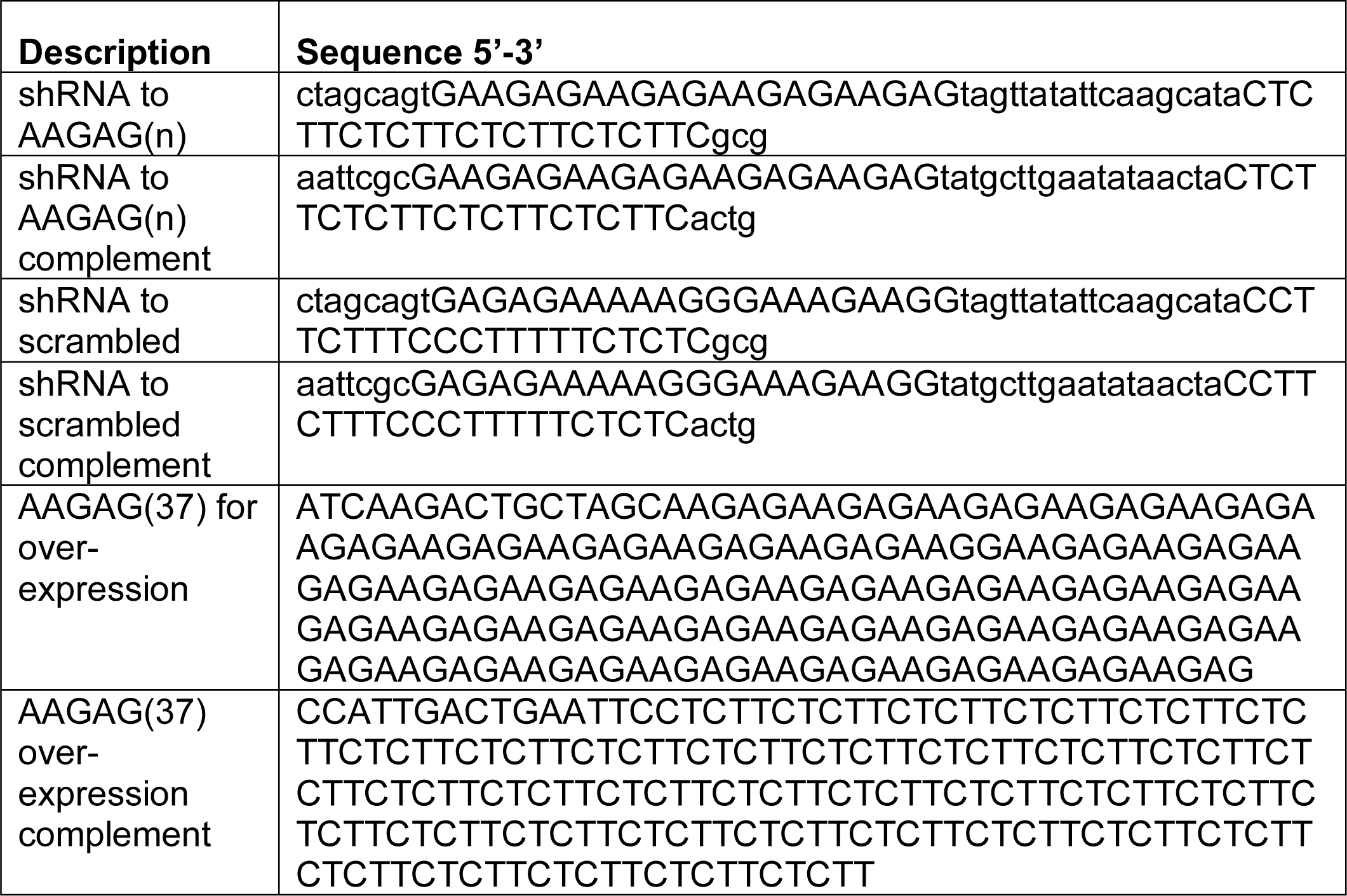
shRNA oligos and overexpression oligos.

### Viability assay

y[1] v[1]:UAS-AAGAGshRNA:: (shRNA to AAGAG), y[1] v[1]:UAS-scramble shRNA:: (shRNA to scrambled) y[1] sc[*] v[1]; P{y[+t7.7] v[+t1.8]=VALIUM20-mCherry}attP2 (dsRNA to mCherry) males were crossed to y[1] w[*]:: P{w[+mC]=Act5C-GAL4}17bFO1/TM6B, Tb[1] (actin-GAL4/Tubby) female virgins. For calculation of ratios of RNAi/Tubby control prior to pupal stage (Extended Data Fig. 6G), the numbers of non-Tubby (RNAi) and Tubby pupae were scored. For each parental cross, a minimum of 11 biological replicates were completed at 25°C, and each vial included at least 8 and no more than 43 pupae of any individual genotype. p-value (2 tailed, type 3): **p=0.013. For calculation of death rates during different stages of development, (Extended Data Fig. 6H), we used the following: To determine L1-L2 death rates, L1 and L2 Tubby and non-Tubby (RNAi) larvae were transferred to separate vials. Those that did not survive to visible L3 were scored as dead. To determine L3 death rates, L3 from lay plates were transferred to vials and those that did not survive to pupae were scored as dead. For pupal lethality, non-eclosed pupae from L1-L2, and L3 transfers were scored as dead. L1-L2 death rate (min. n L1/L2 analyzed per parental set of three experiments = 7 L1/L2): p-values (2 tailed, type 3): A/M= AAGAG to mCherry; A/S= AAGAG to Scrambled; S/M= scrambled to mCherry; A/S=0.457, A/M=0.404; L3 death rate (min. n per 5 parental sets of L3 analyzed=7 L3): A/S=0.125, A/M=0.019; Pupal death rate (min. n per 3 parental sets of pupae analyzed = 10 pupae): A/S=0.002, A/M=0.992, S/M=0.002. Of note, the high pupal death in scrambled control is perplexing considering that we could not find mRNAs that would be targeted by this hairpin. We speculate that this lethality results from off-target effects on un-annotated RNA, and/or the hairpin RNA is toxic. Importantly, however, the lethal phase differed between AAGAG RNAi (L1-L3) vs scrambled RNAi (pupal) (Extended Data Fig. 6H).

**Table 4.** Antibodies used for Immuno-Fluorescence.

| antibody | Supplier; Cat. number | working concentration |
| --- | --- | --- |
| Rabbit-anti H3K9me3 | Abcam; 8898 | 1/250 |
| Mouse-anti H3K9me2 | Active Motif; 39753 | 1/250 |
| Rabbit-anti-H2AV | Lake placid AM318; 9751 | 1/100 |
| Goat anti-GFP | Rockland 121600-101-215 | 1/500 |
| Rabbit anti-H4acetyl | Millipore 06-598 | 1/200 |
| Rat anti-Mst77F | Elaine Dunleavy, PhD; NUI Galway, Ireland | 1/200 |
| Guinea pig anti-Mst35Ba/Bb (Protamine A/B) | Elaine Dunleavy, PhD; NUI Galway, Ireland | 1/200 |

**Table 5.** Fly lines.

| Stock name or genotype | Obtained from: stock number | Description |
| --- | --- | --- |
| y[1] v[1]; P{y[+t7.7]=CaryP}attP40 | Bloomington: 36304 | Background strain for insertion of pValium20 vector containing shRNA |
| y[1] v[1]; P{y[+t7.7]=CaryP}attP2 | Bloomington:<br>36303 | Background strain for insertion of pValium vector containing AAGAG expression construct. |
| y[1] sc[*] v[1]; P{y[+t7.7] v[+t1.8]=VALIUM20-mCherry}attP2 | Bloomington:<br>35785 | Control strain for RNAi. Expresses dsRNA to mCherry |
| y[1] v[1]:UAS-AAGAG shRNA:: | Rainbow Transgenic Flies, Inc | Expresses shRNA under UAS promoter targeting AAGAG(n) |
| y[1] v[1]:UAS-scramble shRNA:: | Rainbow Transgenic Flies, Inc | Expresses shRNA under UAS promoter targeting random AG containing sequences |
| y[1] w[67c23]; P{w[+mC]=dpp-GAL4.PS}6A/TM3, Ser[1] | Bloomington:<br>7007 | Dpp-GAL4 |
| y[1] v[1]:UAS-AAGAG(37) | Rainbow Transgenic Flies, Inc | Expresses a 187 base repeat of AAGAG RNA under a UAS promoter |
| <u>C(1;Y)1, y[1] w[A738]: y[+]/0 & C(1)RM, y[1] v[1]/0</u> | Bloomington:24<br>94 | XO (Y chromosome deficient males) |
| y[*] w[*]; P{w[+mW.hs]=GawB}NP1233 / CyO, P{w[-]=UAS-lacZ.UW14}UW14 | <u>Kyoto: 103948</u> | Fascillin-GAL4 |
| y[*] w[*]; P{w[+mW.hs]=GawB}NP1624 / CyO, P{w[-]=UAS-lacZ.UW14}UW14 | <u>Kyoto:104055</u> | Traffic Jam-GAL4 |
| w[*]; P{w[+mW.hs]=GawB}ptc[559.1] | <u>kyoto: 103948</u> | PTC-GAL4 |
| :: nanos-Gal4, dcr2-UAS / TM3 sb | Unknown | Nanos-GAL4 |
| w;;bamGAL4, UAS-dicer2 | Unknown | Bam-GAL4 |
| y[1] w[*]: P{w[+mC]=Act5C-GAL4}17bFO1/TM6B, Tb[1] | Bloomington:<br>stock 3954 | Expresses GAL4 ubiquitously under control of Act5C promoter |

### Fertility assay

Flies containing shRNA to AAGAG or scrambled control were mated to different testes GAL4 drivers (Extended Data Table 2) at 25°C, in at least duplicate parental (F_0_) sets. From each parental set, individual F_1_ male progeny (minimum of 12 per parental set) were then allowed to mate with two female Oregon R virgins for 10 days at 25°C. Male flies were counted as sterile if, after 10 days, the male and at least one female were still alive and no larvae, pupae or adult F_2_ progeny present. Female fertility was calculated as above, with one female RNAi and two Oregon R males. For Bam-GAL4 driven RNAi, female fertility was calculated as above from a minimum of three parental (F_0_) sets using a minimum of 10 F_1_ progeny for each. Scrambled RNAi male fertility for this cross was calculated as above from a minimum of four parental (F_0_) sets, using a minimum of 11 F_1_ progeny. AAGAG Bam-GAL4 driven RNAi male fertility of 0% was calculated from >>10 (F_0_) parental sets, hundreds of F_1_ individual males, and at both 25°C and 29°C. For rescue experiments, triplicate parental sets were used, where one F_1_ male (minimum 15 per parental set) was mated to three Oregon R virgin females for 10 days and fertility assayed as above.

### Morphology defects in RNAi sperm

For quantification of abnormalities in sperm DNA morphology (Fig. 3E), a minimum of four testes, each from a different male, were analyzed per genotype (see Table 6 below). Essentially, a projection image of the basal end of testes was made using a 40x confocal objective and all sperm DNA bundles were scored. See Fig. 3B for examples of sperm DNA morphology.

**Table 6.** Quantification of sperm DNA morphological defects in AAGAG and scrambled RNAi 4-7 day old testes.

|  | Testes | Normal Bundle | Lagging bundle | Kinked | Knotted | Needle Eyed | Decondensed |
| --- | --- | --- | --- | --- | --- | --- | --- |
| **AAGAG RNAi** | 1 | 0 | 8 | 2 | 0 | 0 | 2 |
|  | 2 | 1 | 8 | 0 | 2 | 2 | 0 |
|  | 3 | 0 | 1 | 5 | 2 | 1 | 0 |
|  | 4 | 0 | 0 | 1 | 0 | 0 | 0 |
|  | 5 | 0 | 1 | 2 | 1 | 1 | 0 |
|  | 6 | 0 | 1 | 5 | 1 | 0 | 0 |
|  | 7 | 0 | 0 | 3 | 3 | 0 | 0 |
|  | 8 | 0 | 0 | 0 | 0 | 1 | 0 |
| **Scrambled RNAi** | 1 | 2 | 2 | 4 | 0 | 0 | 0 |
|  | 2 | 9 | 7 | 6 | 0 | 0 | 0 |
|  | 3 | 21 | 5 | 0 | 0 | 0 | 0 |
|  | 4 | 5 | 1 | 0 | 0 | 0 | 0 |

### Cross to make XO males

For analysis of AAGAG RNA levels in male testes without a Y-chromosome, y[1]w[1] males were mated to C(1)RM, y[1] v[1]/0 females (Bloomington stock # 2494) at 25°C and 0-6hr testes from F_1_ males imaged.

### qPCR conditions

RNA was extracted and cDNA made by established methods. For qPCR, 10µl 2X Absolute Blue qPCR SYBR low Rox mix (Thermofisher, cat. AB4318) was added, forward and reverse oligos each to 0.15µM, 0.5µl cDNA, and water to 20µl. qPCR conditions were as follows: 95°C, 15min; 40cycles (95°C 15s, 58°C 30s., 72°C 30s); 72°C 30sec performed on AB 7500 Fast Real Time PCR System.

**Table 7.**
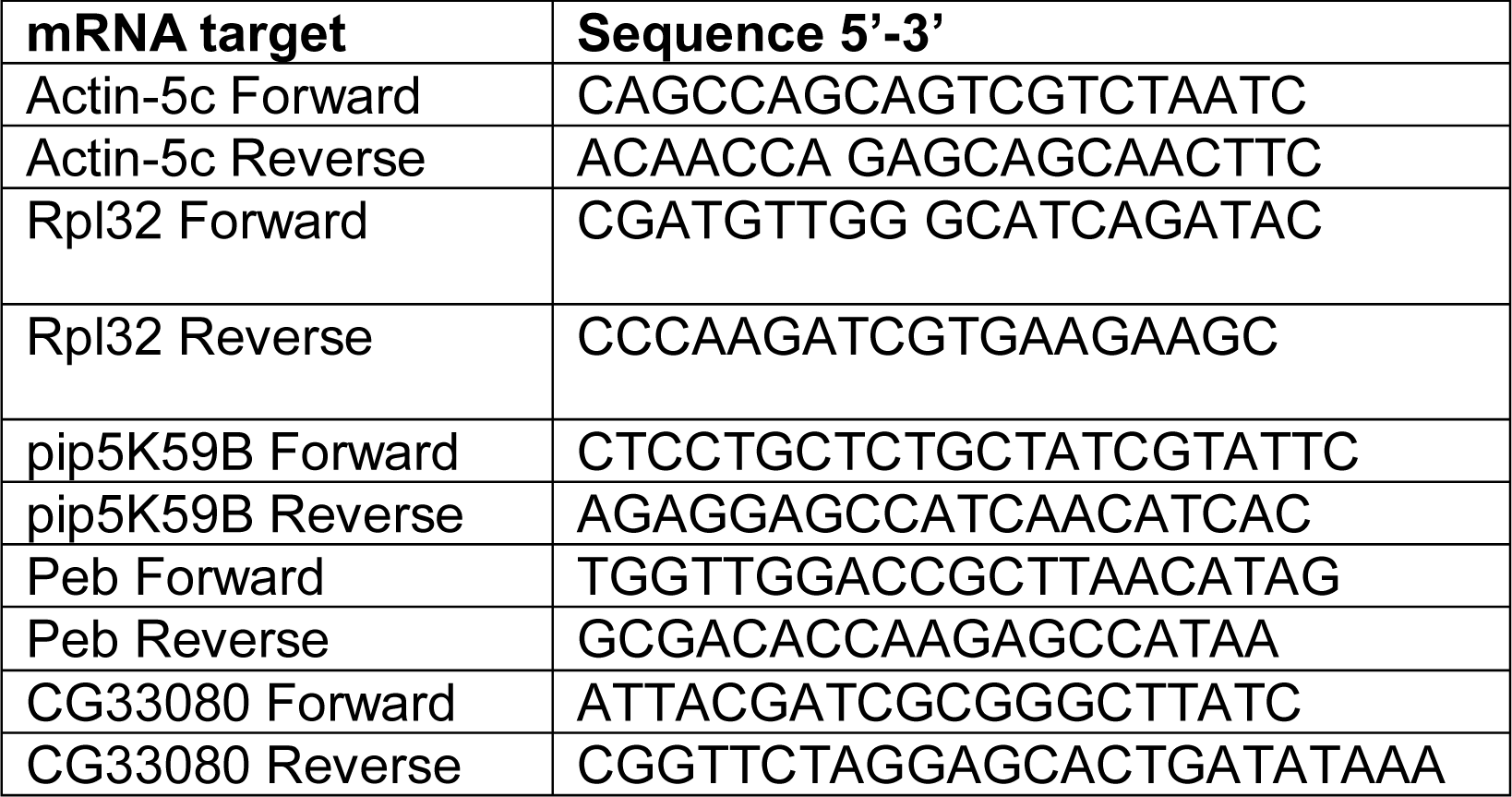
qPCR oligos.

## Bibliography

1. Sun, X., Le, H. D., Wahlstrom, J. M. & Karpen, G. H. Sequence Analysis of a Functional Drosophila Centromere. Genome Res. 13, 182–194 (2003).

2. Hoskins, R. A. et al. Sequence finishing and mapping of Drosophila melanogaster heterochromatin. Science 316, 1625–1628 (2007).

3. Dernburg, A. F., Sedat, J. W. & Hawley, R. S. Direct Evidence of a Role for Heterochromatin in Meiotic Chromosome Segregation. Cell 86, 135–146 (1996).

4. Ferree, P. M. & Barbash, D. A. Species-Specific Heterochromatin Prevents Mitotic Chromosome Segregation to Cause Hybrid Lethality in Drosophila. PLoS Biol 7, e1000234–13 (2009).

5. Yap, K. et al. A Short Tandem Repeat-Enriched RNA Assembles a Nuclear Compartment to Control Alternative Splicing and Promote Cell Survival. Molecular Cell 72, 525–540.e13 (2018).

6. Jain, A. & Vale, R. D. RNA phase transitions in repeat expansion disorders. Nature Publishing Group 546, 243–247 (2017).

7. Zhu, Q. et al. BRCA1 tumour suppression occurs via heterochromatin-mediated silencing. Nature Publishing Group 477, 179–184 (2011).

8. McNulty, S. M., Sullivan, L. L. & Sullivan, B. A. Human Centromeres Produce Chromosome-Specific and Array-Specific Alpha Satellite Transcripts that Are Complexed with CENP-A and CENP-C. Developmental Cell 42, 226–240.e6 (2017).

9. Johnson, W. L. et al. RNA-dependent stabilization of SUV39H1 at constitutive heterochromatin. eLife 6, e25299–32 (2017).

10. Camacho, O. V. et al. Major satellite repeat RNA stabilize heterochromatin retention of Suv39h enzymes by RNA-nucleosome association and RNA:DNA hybrid formation. eLife 6, e25293–29 (2017).

11. Shirai, A. et al. Impact of nucleic acid and methylated H3K9 binding activities of Suv39h1 on its heterochromatin assembly. eLife 6, e25317–23 (2017).

12. Rošić, S., Köhler, F. & Erhardt, S. Repetitive centromeric satellite RNA is essential for kinetochore formation and cell division. J Cell Biol 207, 335–349 (2014).

13. Lohe, A. R., Hilliker, A. J. & Roberts, P. A. Mapping simple repeated DNA sequences in heterochromatin of Drosophila melanogaster. Genetics 134, 1149–1174 (1993).

14. Rinn, J. L. & Chang, H. Y. Genome Regulation by Long Noncoding RNAs. Annu. Rev. Biochem. 81, 145–166 (2012).

15. Brown, J. B. et al. Diversity and dynamics of the Drosophila transcriptome. Nature Publishing Group 1–7 (2014). doi:10.1038/nature12962

16. Pathak, R. et al. AAGAG repeat RNA is an essential component of nuclear matrix in Drosophila. RNA Biology 10, 564–571 (2014).

17. Strom, A. R. et al. Phase separation drives heterochromatin domain formation. Nature Publishing Group 547, 241–245 (2017).

18. Yuan, K. & O’Farrell, P. H. TALE-light imaging reveals maternally guided, H3K9me2/3-independent emergence of functional heterochromatin in Drosophila embryos. Genes & Development 30, 579–593 (2016).

19. White-Cooper, H. Tissue, cell type and stage-specific ectopic gene expression and RNAi induction in the Drosophila testis. Spermatogenesis 2, 11–22 (2012).

20. Rathke, C. et al. Distinct functions of Mst77F and protamines in nuclear shaping and chromatin condensation during Drosophila spermiogenesis. European Journal of Cell Biology 89, 326–338 (2010).

21. Raja, S. J. & Renkawitz-Pohl, R. Replacement by Drosophila melanogaster Protamines and Mst77F of Histones during Chromatin Condensation in Late Spermatids and Role of Sesame in the Removal of These Proteins from the Male Pronucleus. Molecular and Cellular Biology 26, 3682–3682 (2006).

22. Wei, K. H. C. et al. Variable Rates of Simple Satellite Gains across the Drosophila Phylogeny. Molecular Biology and Evolution 35, 925–941 (2018).

23. Lohe, A. R. & Brutlag, D. L. Multiplicity of satellite DNA sequences in Drosophila melanogaster. Proceedings of the National Academy of Sciences 83, 696–700 (1986).

24. Chen, S., Krinsky, B. H. & Long, M. New genes as drivers of phenotypic evolution. Nature Reviews Genetics 14, 645 EP ––660 (2013).

25. Ding, Y. et al. A Young Drosophila Duplicate Gene Plays Essential Roles in Spermatogenesis by Regulating Several Y-Linked Male Fertility Genes. PLoS Genet 6, e1001255–12 (2010).

26. Kaessmann, H. Origins, evolution, and phenotypic impact of new genes. Genome Res. 20, 1313–1326 (2010).

27. Legendre, F. et al. Whole Mount RNA Fluorescent *in situ* Hybridization of *Drosophila* Embryos. JoVE 1–8 (2013). doi:10.3791/50057

28. Shpiz, S., Lavrov, S. & Kalmykova, A. in Programmed Cell Death 1093, 161–169 (Humana Press, 2013).

29. Jandura, A., Hu, J., Wilk, R. & Krause, H. M. High Resolution Fluorescent In Situ Hybridization in Drosophila Embryos and Tissues Using Tyramide Signal Amplification. JoVE e56281–e56281 (2017). doi:10.3791/56281

30. Cai, W., Jin, Y., Girton, J., Johansen, J. & Johansen, K. M. Preparation of Drosophila polytene chromosome squashes for antibody labeling. JoVE e1748–e1748 (2010). doi:10.3791/1748

31. Nicholson, A. W. Ribonuclease III mechanisms of double-stranded RNA cleavage. Wiley Interdiscip Rev RNA 5, 31–48 (2014).

32. Herbert, C., Dzowo, Y. K., Urban, A., Kiggins, C. N. & Resendiz, M. J. E. Reactivity and Specificity of RNase T1, RNase A, and RNase H toward Oligonucleotides of RNA Containing 8-Oxo-7,8-dihydroguanosine. Biochemistry 57, 2971–2983 (2018).

33. Kelemen, B. R., Schultz, L. W., Sweeney, R. Y. & Raines, R. T. Excavating an active site: the nucleobase specificity of ribonuclease A. Biochemistry 39, 14487–14494 (2000).

34. Hoskins, R. A. et al. The Release 6 reference sequence of the Drosophila melanogaster genome. Genome Res. 25, 445–458 (2015).

35. Ni, J.-Q. et al. A genome-scale shRNA resource for transgenic RNAi in Drosophila. Nat Meth 8, 405–407 (2011).

