## Supplementary material for "RNA transcribed from heterochromatic simple-tandem repeats are required for male fertility and histone-protamine exchange in *Drosophila melanogaster*"

a

| Repeat | RNA-Foci? | Repeat | RNA-Foci? |
|---|---|---|---|
| AAACC(n) | ND | AGCC(n) | ND |
| AAACG(n) | ND | AGCCCC(n) | ND |
| AAAG(n) | ND | AGCCG(n) | ND |
| AACAG(n) | ND | AGCGG(n) | ND |
| AACGG(n) | ND | AGTCT(n) | ND |
| AACTC(n) | ND | ATATC(n) | ND |
| AAGAGG(n) | ND | ATC(n) | ND |
| AAGCG(n) | ND | ATCGC(n) | ND |
| AAGGAG(n) | ND | ATTAC(n)* | ND |
| AATAG(n) | ND | dodeca | ND |
| AATAGAC(n) | ND | AACAC(n)* | + |
| AATC(n) | ND | AATAC(n)* | + |
| AATT(n) | ND | 359 satellite* | + |
| ACCC(n) | ND | AATAACATAG(n) | + |
| ACCCC(n) | ND | AAGAG(n)* | + |
| AGC(n) | ND | | |

b

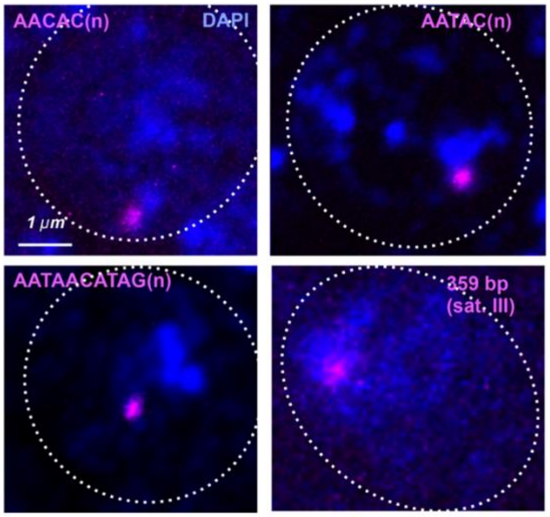


**Extended Data Figure 1. RNA-FISH analysis of satellite RNAs in cycle 14 embryos. a,** ND=Not detected, *repeats tested for expression from both strands. For more details about AAGAG RNA in cycle 14, see Figs. 1A and C and Extended Data Fig. 2. **b,** Projections of cycle 14 nuclei (dashed circle = nuclear periphery); DNA (DAPI)= blue, indicated satellite RNAs in magenta.

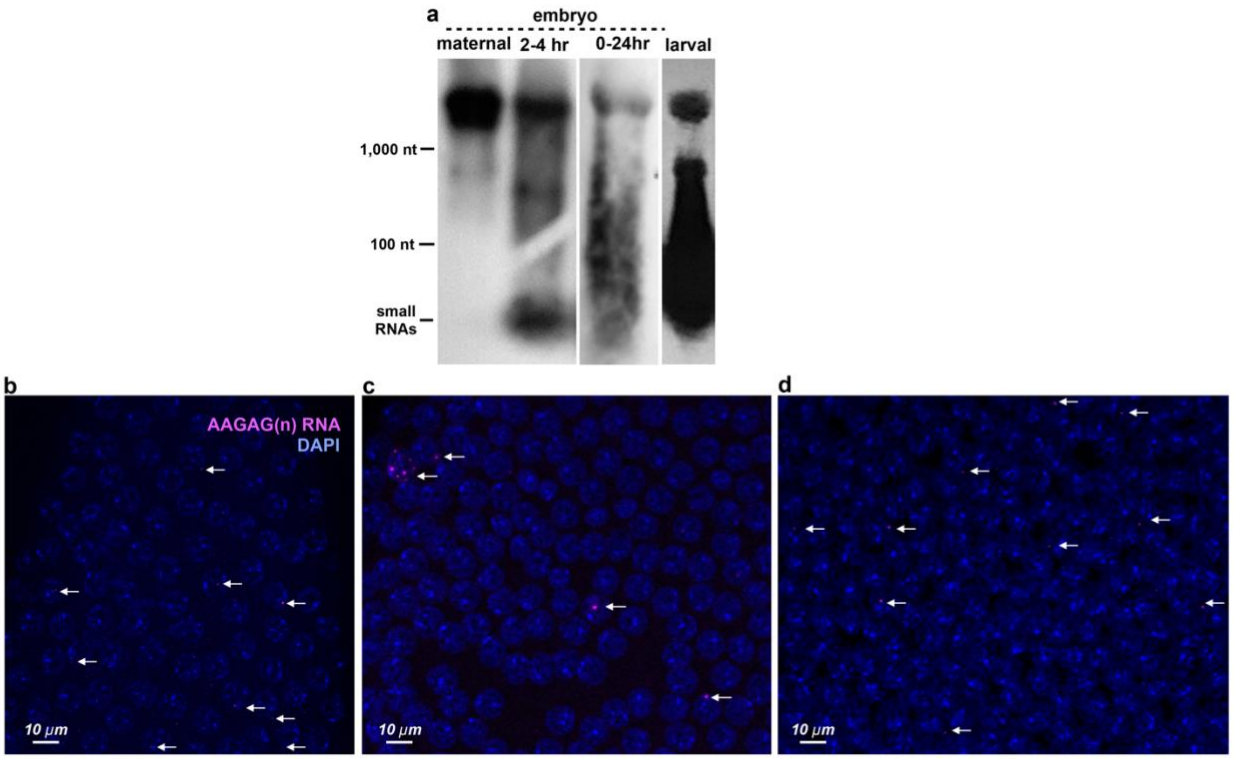


**Extended Data Figure 2.** **a,** Northern blot analyzing RNA from Oregon R embryos and third instar larva hybridized with a probe complementary to AAGAG(n). Note that signal intensity is not a representation of relative AAGAG RNA levels since different exposure times were used. **b-d,** Examples of AAGAG RNA distributions in cycle 12, 13 and 14 embryos. Projections through embryo nuclei stained with DAPI (Blue). AAGAG RNA foci are shown in magenta and marked with arrows. **b**, One of the 33% of cycle 12 embryos with AAGAG RNA foci. **c**, One of the 67% of cycle 13 embryos with one or more foci **d,** Cycle 14 embryo.

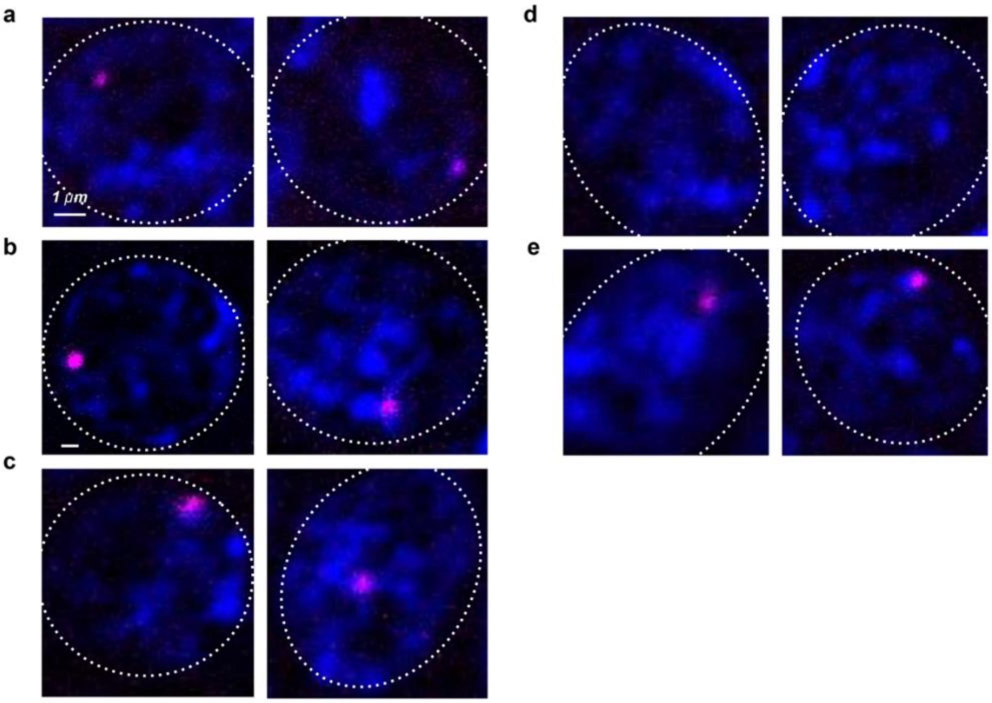


**Extended Data Figure 3. AAGAG RNA foci contain single-stranded RNA and are not associated with R-loops.** Confocal sections of embryonic nuclei in cycle 14 (with exception of left panel in ‘b’), nuclear periphery outlined in dotted circles. **a**, No RNase control. **b**, Treated with RNaseIII (left nucleus is cycle 12) **c**, RNaseH **d**, RNase1 and **e**, RNaseA

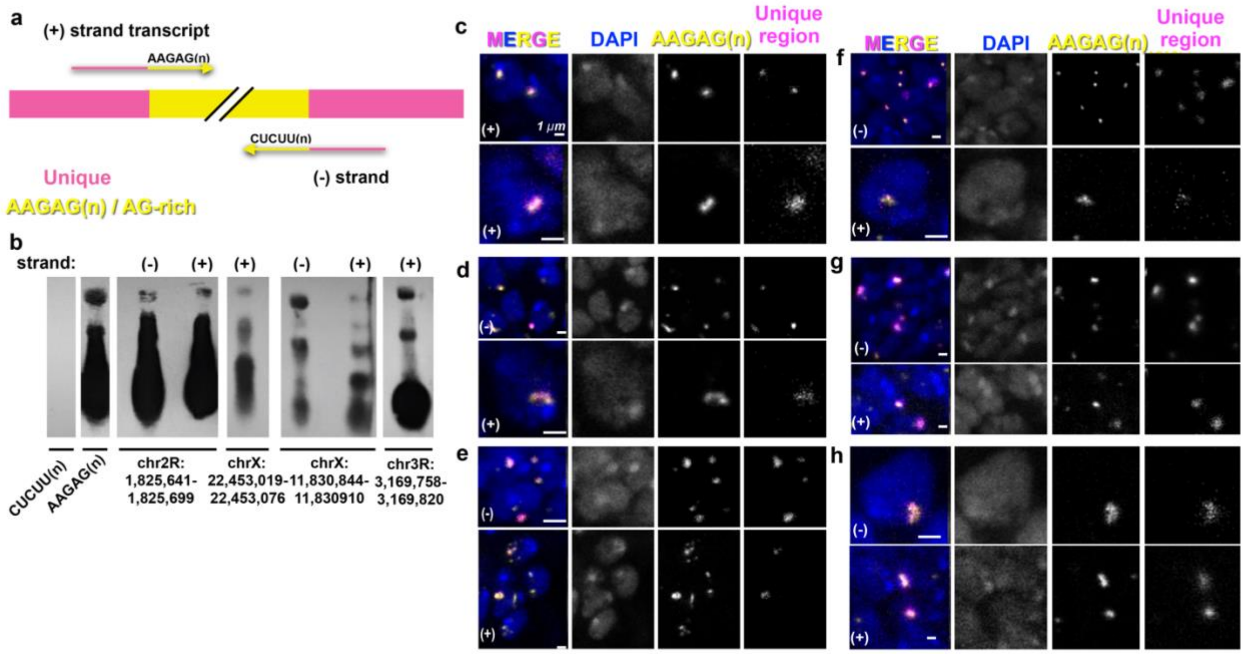


**Extended Data Figure 4. AAGAG RNA transcripts originate from 2R, X and 3R heterochromatin loci and are transcribed in embryos and larval brain. a**, Unique regions adjacent to AAGAG(n), or AAGAG(n) within AG rich regions, were identified as potential sources of satellite transcripts, as described in Materials and Methods. (+) indicates transcript containing AAGAG(n) or AG(n) blocks, while (-) indicates transcript containing CUCUU(n) or CU(n) blocks **b**, Northern blots of L3 RNA using probes to satellite (AAGAG or CUCUU), or adjacent unique regions. Unique regions shown are those containing at least one similar band size as AAGAG RNA. **c**—**h**, Confocal sections of embryo ventral ganglia or L3 brain lobe nuclei stained with DAPI (blue), and RNA-FISH to AAGAG (yellow) and unique region locations (magenta). ‘Unique Region’ (single copy sequence) RNA-FISH required Tyramide Signal Amplification (TSA), and therefore displays poorer resolution compared to AAGAG RNA (detected without TSA). Images labelled (+) used probes complementary to the strand containing AAGAG(n) or AG(n) blocks, while those labelled (-) recognize the strand containing CUCUU(n) or CU(n) blocks. Note that ‘unique region’ probe binds to regions adjacent to AAGAG(n) or AG(n), and not AAGAG(n) or AG(n) sequences themselves. Also note that the chr3R region indicated in ‘b’ was not analyzed in larvae. **c**, Nuclei from late embryo ventral ganglia, RNA-FISH to AAGAG and probe from genomic coordinates chr2R:1,825,641-1,825,699 (top) or chrX:12,660,096-12,660,145 (bottom). **d**-**h,** Confocal sections from L3 brain lobe nuclei, RNA-FISH to AAGAG and the following: **d,** Unique region chr2R:1, 825,641-1825699 **e,** chr2R:1,826,691-1,826,740 **f**, chrX:11,830,844-11,830910 **g**, chrX:12,660,096-12,660,145 **h**, chrX:22,453,019-22,453,076

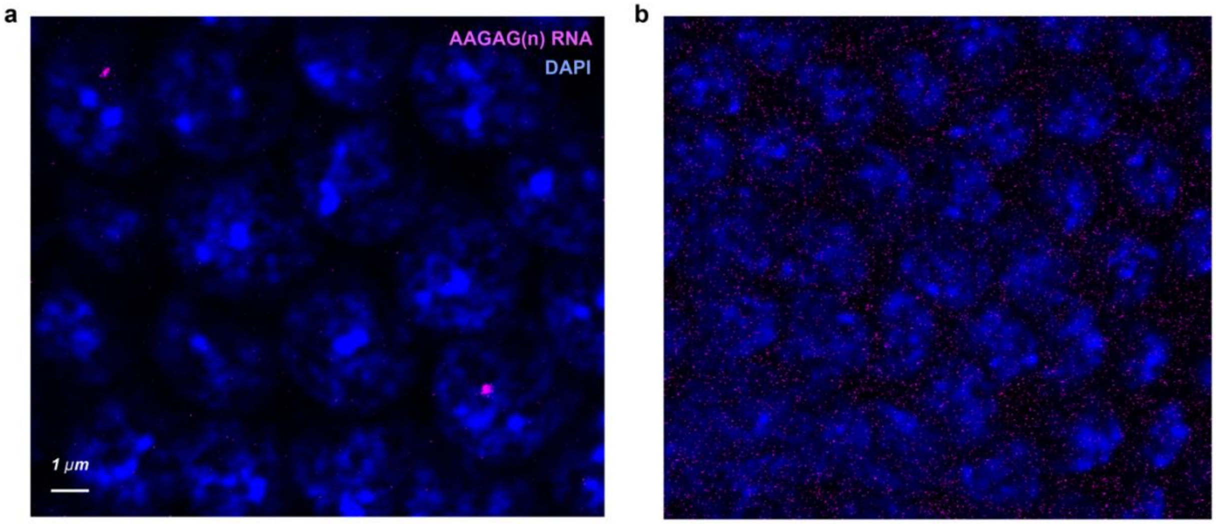


**Extended Data Figure 5. AAGAG RNA-FISH localizes RNA and not DNA.** Confocal sections of cycle 14 nuclei treated with either **a**, RNAseH or **b**, RNAseIII after AAGAG RNA probe (magenta) hybridization. A higher laser intensity for the AAGAG probe channel was used in **b** to demonstrate abolishment of AAGAG signal.

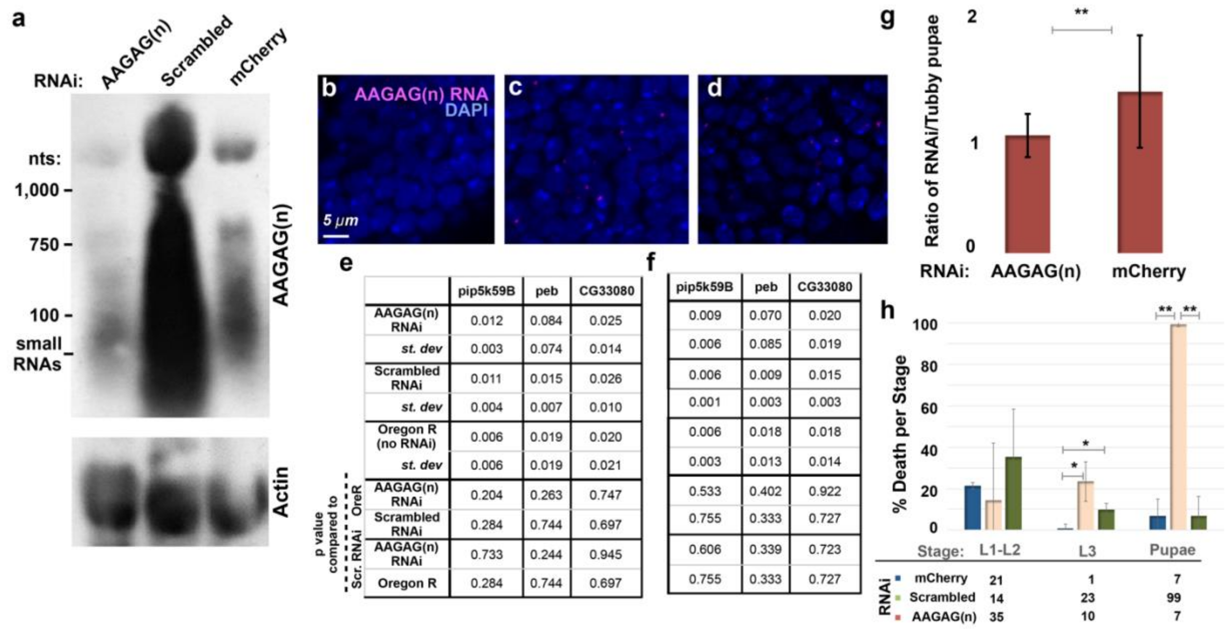


**e**

| | pip5k59B | peb | CG33080 |
|---|---|---|---|
| AAGAG(n) RNAi | 0.012 | 0.084 | 0.025 |
| *st. dev* | 0.003 | 0.074 | 0.014 |
| Scrambled RNAi | 0.011 | 0.015 | 0.026 |
| *st. dev* | 0.004 | 0.007 | 0.010 |
| Oregon R (no RNAi) | 0.006 | 0.019 | 0.020 |
| *st. dev* | 0.006 | 0.019 | 0.021 |
| p value compared to OreR: AAGAG(n) RNAi | 0.204 | 0.263 | 0.747 |
| p value compared to OreR: Scrambled RNAi | 0.284 | 0.744 | 0.697 |
| p value compared to Scr. RNAi: AAGAG(n) RNAi | 0.733 | 0.244 | 0.945 |
| p value compared to Scr. RNAi: Oregon R | 0.284 | 0.744 | 0.697 |

**f**

| | pip5k59B | peb | CG33080 |
|---|---|---|---|
| AAGAG(n) RNAi | 0.009 | 0.070 | 0.020 |
| *st. dev* | 0.006 | 0.085 | 0.019 |
| Scrambled RNAi | 0.006 | 0.009 | 0.015 |
| *st. dev* | 0.001 | 0.003 | 0.003 |
| Oregon R (no RNAi) | 0.006 | 0.018 | 0.018 |
| *st. dev* | 0.003 | 0.013 | 0.014 |
| p value compared to OreR: AAGAG(n) RNAi | 0.533 | 0.402 | 0.922 |
| p value compared to OreR: Scrambled RNAi | 0.755 | 0.333 | 0.727 |
| p value compared to Scr. RNAi: AAGAG(n) RNAi | 0.606 | 0.339 | 0.723 |
| p value compared to Scr. RNAi: Oregon R | 0.755 | 0.333 | 0.727 |

**Extended Data Figure 6. AAGAG RNA is decreased and foci abolished in L3 with actin-GAL4 driven RNAi to AAGAG, without affecting levels of genes whose mRNAs contain short runs of AAGAG. Also, AAGAG RNAi results in lethality. a,** Northern blot with probes to AAGAG RNA or actin-5c in L3 RNAi. The AAGAG RNAi L3 AAGAG RNA top band signal is approximately 86% and 75% reduced compared to either scrambled or mCherry controls, respectively, when normalized to the actin-5c loading control. **b** and **c,** Confocal sections of brain lobes stained with DAPI (blue) and RNA-FISH to AAGAG RNA (magenta) imaged with the same intensity. **b,** Brain lobe from AAGAG RNAi **c,** Brain lobe from scrambled RNAi. **d**, Brain lobe from mCherry RNAi. **e,** RNA transcript levels (qRT-PCR) for euchromatic genes whose mRNAs contain short runs of AAGAG (pip5k59B, peb, CG33080). Numbers are means (from three biological replicates) ± standard deviation, after normalization to either **e**, actin-5c loading control, or **f**, rpl32 loading control. t-tests were performed in comparison to Oregon R or scrambled RNAi controls. This demonstrates that RNA levels of the few mRNAs containing an AAGAG sequence are not affected by AAGAG RNAi, ruling out the possibility that the observed lethality is due to off-target effects. **g,** Ratio of pupae containing RNAi (driven by ubiquitous actin-GAL4 driver) or Tubby control, demonstrating lethality in AAGAG RNAi prior to the adult stage. **h,** For embryos that hatch, death rates in larval and pupal stages differ after RNAi depletion of AAGAG, scrambled and mCherry controls (driven by ubiquitous actin-GAL4 driver). Note that death rate per stage is a measure of death only for those that survive to the indicated stage. $^{**}p \leq 0.01$, $^{*}p \leq 0.05$; error bars = SD.

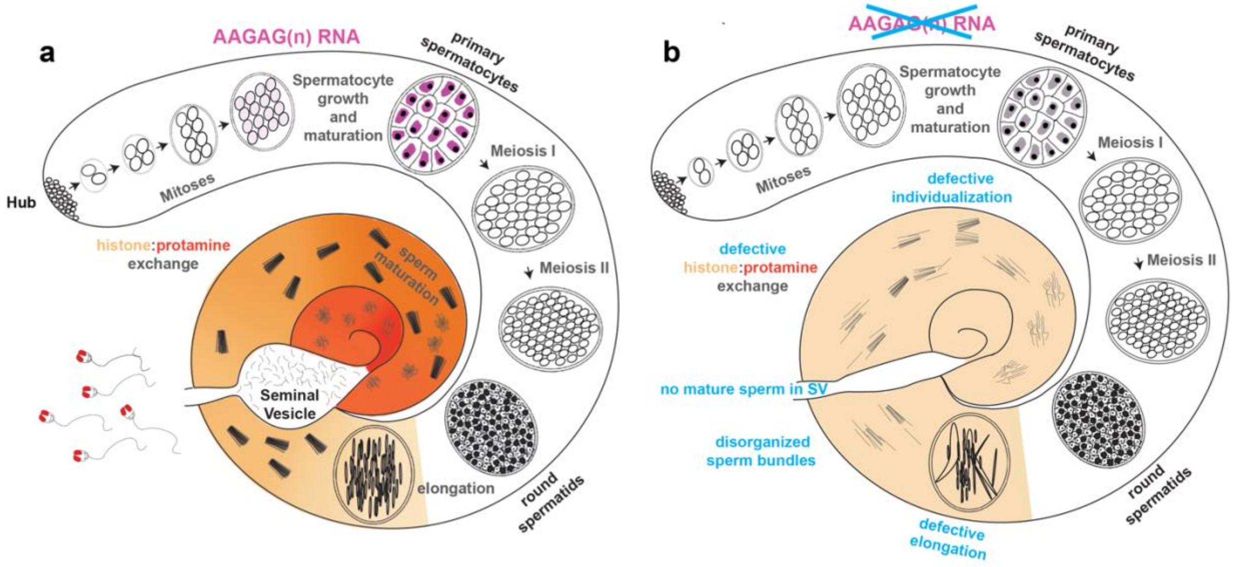


**Extended Data Figure 7. Overview of normal spermatogenesis and defects observed after AAGAG RNA depletion. a,** Spermatogenesis in *Drosophila melanogaster* initiates at the apical end of the testes (Hub), where GSCs divide asymmetrically, producing gonialblasts (GBs) that begin cell-differentiation. GB cells then undergo four mitotic divisions with incomplete cytokinesis to produce a cyst of 16 primary spermatocytes. Spermatocytes then undergo pre-meiotic S phase, mature during a prolonged G2 phase, and increase substantially in volume. The majority of testes-specific gene expression occurs at the primary spermatocyte stage, while genes not required until later stages are translationally repressed. (reviewed in [1]). Mature spermatocytes then undergo two rounds of meiosis to produce round spermatids[2], which are then processed into independent, condensed sperm nuclei in two stages[3-5]. First, round spermatids undergo chromatin compaction, acrosome formation and flagellar elongation[3,4]. During chromatin compaction, a wave of histone H4 acetylation occurs, followed by deposition of the transition protein Mst77f[6]. Next, transition proteins are removed followed by the incorporation of protamines and prtl99c (histone:protamine exchange, indicated by tan to deep orange gradient)[3,4]. Finally, spermatid individualization involves removal of cytoplasm and tight condensing and coiling of chromatin[5]. Mature sperm are then stored in the seminal vesicle. **b,** Summary of defects in late stages of spermatogenesis observed after depletion of AAGAG RNA by RNAi, using the Bam-Gal4 driver (data in Figure 3). Although AAGAG RNA is not visible in normal testes after the S6 spermatocyte stage (see **a**), RNAi depletion of AAGAG RNA only produces visible defects after the round spermatid stage. Aberrant elongation, sperm bundles, and defective histone:protamine exchange likely cause the observed complete absence of mature sperm in the SV.

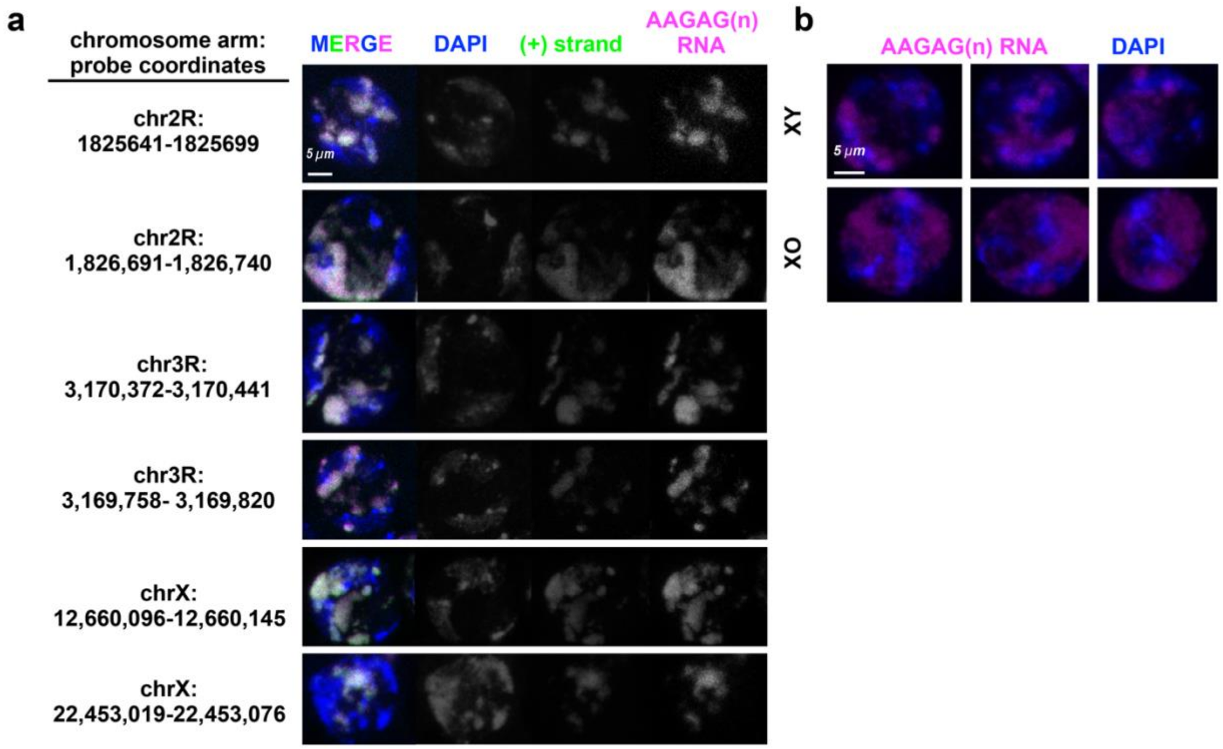


**Extended Data Figure 8. Heterochromatic regions adjacent to AAGAG(n) or AG(n)-rich blocks are transcribed in primary spermatocytes, co-localize with AAGAG(n) RNA foci and do not come from the Y. a,** Projections of Oregon R S5 spermatocytes probed for unique regions of RNA (green) adjacent to AAGAG(n) (magenta) or AAGAG(n) containing AG rich blocks. DAPI (DNA) is indicated in blue. **b**, Projections of S5 spermatocyte probed to AAGAG RNA (magenta) imaged at same laser intensities in XY and XO genotypes. DNA is stained with DAPI (blue).

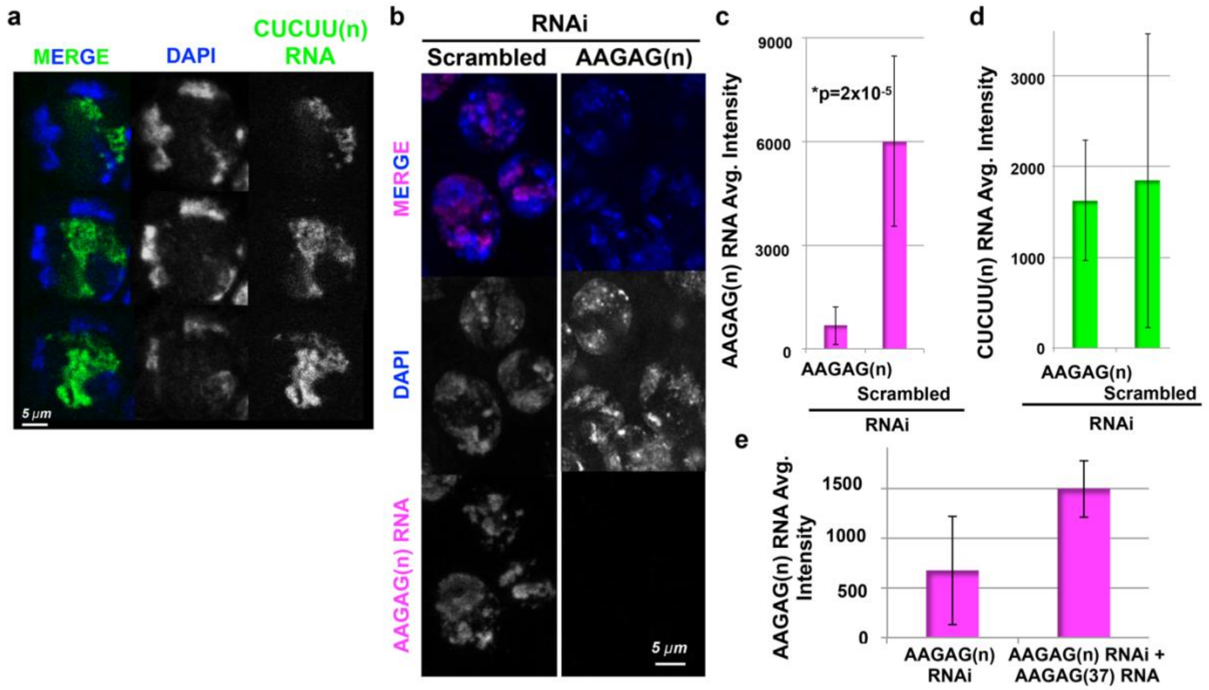


**Extended Data Figure 9. AAGAG RNA and not CUCUU RNA is substantially decreased in Bam-GAL4 driven AAGAG RNAi, and AAGAG RNA levels are increased in rescue experiments a,** Although absent in embryos and somatic larval tissues, CUCUU RNA (green) is expressed in adult spermatocytes. Note that CUCUU RNA is localized to the S5 lumen, internal to the chromatin (DAPI), in contrast to the peripheral localization of AAGAG RNA (see Figure 3b); DNA = DAPI (blue). **b,** Projections of AAGAG foci (magenta) in S5 spermatocytes after Bam-GAL4 driven Scrambled control or AAGAG RNAi. Signal was imaged with the same laser intensities for each genotype. **c** and **d**, Average median intensities (arbitrary units, ± st. dev.) of **c**, AAGAG RNA, $p=2x10^{-5}$ and **d**, CUCUU RNA in S5 spermatocytes in AAGAG and scrambled RNAi testes (not significant). This represents a 72% reduction of AAGAG RNA in S5 spermatocytes after AAGAG RNAi, compared to scrambled controls, with little to no decrease in CUCUU RNA. **e,** Average intensity of AAGAG RNA in S5 spermatocytes after AAGAG RNAi increases significantly (p=0.03) upon co-expression of AAGAG(37) RNA (also induced by the Bam-Gal4 driver).

**Extended Data Table 2: Male fertility in AAGAG RNAi with testes GAL4 drivers upstream of Bam**

| Germline GAL4 RNAi driver | Expression location[7] | % fertile | +/- stdev. | Minimum number of males per set |
|---|---|---|---|---|
| Fascillin | Hub | 94 | 16 | 15 |
| PTC | Soma- CySCs and cyst cells | 90 | 5 | 18 |
| Traffic Jam | Soma- Hub and CySCs | 97 | 4 | 12 |
| Dpp1 | Soma- CySCs and early cyst cells | 96 | 6 | 17 |
| Nanos | Germline- GSCs and early germline cysts | 83 | 5 | 13 |

### Bibliography


1. White-Cooper, H. Molecular mechanisms of gene regulation during Drosophila spermatogenesis. *Reproduction* **139,** 11–21 (2010).
2. McKee, B. D., Yan, R. & Tsai, J.-H. Meiosis in male Drosophila. *Spermatogenesis* **2,** 167–184 (2014).
3. Rathke, C., Baarends, W. M., Awe, S. & Renkawitz-Pohl, R. Chromatin dynamics during spermiogenesis. *Biochim. Biophys. Acta* **1839,** 155–168 (2014).
4. Eren-Ghiani, Z., Rathke, C., Theofel, I. & Renkawitz-Pohl, R. Prtl99C Acts Together with Protamines and Safeguards Male Fertility in Drosophila. *CellReports* **13,** 2327–2335 (2015).
5. Steinhauer, J. Separating from the pack: Molecular mechanisms of Drosophila spermatid individualization. *Spermatogenesis* **5,** e1041345 (2015).
6. Kost, N. *et al.* Multimerization of Drosophila sperm protein Mst77F causes a unique condensed chromatin structure. *Nucleic Acids Research* **43,** 3033–3045 (2015).
7. Demarco, R. S., Eikenes, Å. H., Haglund, K. & Jones, D. L. Investigating spermatogenesis in Drosophila melanogaster. *Methods* **68,** 218–227 (2014).
